# Carriers of mitochondrial DNA macrohaplogroup L3 basic lineages migrated back to Africa from Asia around 70,000 years ago

**DOI:** 10.1101/233502

**Authors:** Vicente M. Cabrera, Patricia Marrero, Khaled K. Abu-Amero, Jose M. Larruga

## Abstract

**Background:** After three decades of mtDNA studies on human evolution the only incontrovertible main result is the African origin of all extant modern humans. In addition, a southern coastal route has been relentlessly imposed to explain the Eurasian colonization of these African pioneers. Based on the age of macrohaplogroup L3, from which all maternal Eurasian and the majority of African lineages originated, that out-of-Africa event has been dated around 60-70 kya. On the opposite side, we have proposed a northern route through Central Asia across the Levant for that expansion. Consistent with the fossil record, we have dated it around 125 kya. To help bridge differences between the molecular and fossil record ages, in this article we assess the possibility that mtDNA macrohaplogroup L3 matured in Eurasia and returned to Africa as basic L3 lineages around 70 kya.

**Results:** The coalescence ages of all Eurasian (M,N) and African L3 lineages, both around 71 kya, are not significantly different. The oldest M and N Eurasian clades are found in southeastern Asia instead near of Africa as expected by the southern route hypothesis. The split of the Y-chromosome composite DE haplogroup is very similar to the age of mtDNA L3. A Eurasian origin and back migration to Africa has been proposed for the African Y-chromosome haplogroup E. Inside Africa, frequency distributions of maternal L3 and paternal E lineages are positively correlated. This correlation is not fully explained by geographic or ethnic affinities. It seems better to be the result of a joint and global replacement of the old autochthonous male and female African lineages by the new Eurasian incomers.

**Conclusions:** These results are congruent with a model proposing an out-of-Africa of early anatomically modern humans around 125 kya. A return to Africa of Eurasian fully modern humans around 70 kya, and a second Eurasian global expansion by 60 kya. Climatic conditions and the presence of Neanderthals played key roles in these human movements.

## BACKGROUND

From a molecular genetics perspective, the hypothesis of a recent African origin of modern humans by around 200 thousand years age (kya) was formulated three decades ago [1]. Today this hypothesis is widely accepted. There is also multidisciplinary agreement that the out-of-Africa expansion of modern humans promoted the extinction of other hominins in Eurasia with only a minor assimilation of their genomes [2]. However, despite the enormous quantity of data accumulated during these years, mainly from the analysis of the mtDNA and Y-chromosome haploid markers, there is a lack of consensus about the time/s and route/s followed by modern humans moving out of Africa. All the indigenous mtDNA diversity outside Africa is comprised into clades M and N, that are branches of the African haplogroup L3 [3–5]. This fact puts a genetic time frame for the out of Africa dispersal around 55-70 kya that is the coalescence age of haplogroup L3 [6]. Likewise, recent Y-chromosome sequence analysis detected a cluster of major non-African founder haplogroups in a short time interval at 47-52 kya [7]. However, it has to be mentioned that the tick of the molecular clock depends on the mutation rate employed [8, 9]. These temporal windows for the exit of modern humans out-of-Africa are at odds with fossil, archaeological and ancient DNA data. Skeletal remains unearthed in the Skhul and Qafzeh caves demonstrated that early modern humans were present in the Levant between 125 and 80 kya [10]. The discovery of modern human teeth in southern China dated to 120-80 kya [11], also supports the presence of anatomically modern humans (AMHS) in eastern Asia during this period. Several archaeological studies uncovered Middle Stone Age (MSA) lithic assemblages, dated around 125-75 kya, in different regions of the Arabian Peninsula, presenting affinities with northeastern African assemblages of the same period [12–14]. These findings suggest that African AMHS may have extended its geographic range to eastern and northern Arabia long before the time frame proposed by molecular data. Ancient DNA (aDNA) analysis is being a major tool in the reconstruction of the past human history. Using these analyses, the Neanderthal introgression into modern humans in Europe has been dated within 35-65 kya [15] which is well in frame with the molecular clock window established for the African exit of modern humans. However, an ancient gene flow from early modern humans into the ancestors of eastern Neanderthals more than 100 kya has been recently reported [16]. These data evidenced that early modern humans and ancestors of Neanderthals from the Siberian Altai region interbred much earlier than previously thought. Besides, whole-genome based studies proposed the split of Eurasian from African populations at 88-112 kya [17], and that the presence of AMHS out of Africa is earlier than 75 kya [18]. A way to avoid all these contradictory pieces of evidence is to state that all these ancient movements out of Africa, prior to 70 kya, did not contribute genetically to present-day human populations. However, we think that major effort should be dedicated to finding a more conciliatory explanation.

Concerning the potential routes followed by modern humans out of Africa, there are two main alternatives not mutually exclusive: a northern dispersal along the Nile-Sinai corridor, and a southern dispersal from the Horn of Africa across the Bab al Mandeb strait. At the beginning of the mtDNA phylogeographic studies, the virtual absence of mtDNA haplogroup M in the Levant, and its presence in Ethiopia, southern Arabia, the Indian subcontinent and East Asia, rendered M the first genetic indicator of a southern route exit from eastern Africa [19]. At that time, it was suggested that this was possibly the only successful early dispersal event of modern humans out of Africa. Shortly after, based on the rarity of mtDNA haplogroup N(xR) in India, and its continuous presence above the Himalayas, we proposed an additional northern route through the Levant [4]. Afterward, intensive and extensive research on mtDNA has been carried out, not only in populations from Central [20–23] and East Asia [24–26] but, also, from the regions covering the hypothetical southern route path as India [27–30], Mainland southeastern Asia [31–33], Island southeast Asia [34–39], New Guinea, North Island Melanesia and Australia [40–44]. The most striking global result was that while in Central and North Asia the indigenous lineages observed were derived branches of southern haplogroups, primary and independent autochthonous M and N clusters were found in every main region of meridional Asia and Australasia. Most surprising was the fact that some N haplogroups in southern China (N10, N11) resulted older than the oldest N western Asia lineages (N1, N2), or that some M haplogroups in Melanesia (M27, Q) were older than the oldest Indian M lineages (M2, M33). Different researchers gave conflicting interpretations to these results. Some perceived them as a confirmation of a rapid southern coastal spread of modern humans from Africa [30, 45–47]. Others claimed an old local population differentiation in each region without any evidence of the shared ancestry expected by the southern dispersal model [48–50]. The existence of a northern route, deduced from the phylogeography of macrohaplogroup N [4], has received additional support from the fossil record [11], from whole genome studies comparing Egyptian and Ethiopian populations [51], and by the fact that all non-African populations present a signal of Neanderthal introgression [52]. However, we realized that what actually macrohaplogroup N points to is a human movement from southeastern Asia to western Asia [53]. We observed the same sense for macrohaplogroup M. In this case, expanding westwards to India [54], and a similar trend follows macrohaplogroup R, the main sister branch of N [55]. Thus, we confirmed that macrohaplogroup M and N indicated, respectively, major southern and northern expansions of modern humans but, ironically, in the opposite sense we predicted long ago [4]. It has to be mentioned that studies based on Y-chromosome sequences also pointed to southeastern Asia as a primitive center of human expansions [56–59]. Now, if we do admit that basic L3 lineages (M, N) have independently evolved in southeastern Asia instead in Africa or near of the African continent borders where the rest of L3 lineages expanded, we are confronted with the dilemma of where the basic trunk of L3 evolved. A gravitating midpoint between eastern Africa and southeastern Asia would situate the born of L3 in inner Asia, with possible opposite expansions back to Africa and forward to eastern Asia. At this point, it has to be mentioned that this possibility has been already modeled, among other options, obtaining the highest likelihood value [60] but, in our opinion, it has not received the attention it deserves. The parallelism of this early back to Africa of mtDNA haplogroup L3 with that proposed for the Y-chromosome haplogroup E [61] is striking.

In this work, we assess the possibility that L3 could have exited from Africa as a pre-L3 lineage that evolved as basic L3 in inner Asia. From there, it came back to Africa and forwarded to southeastern Asia to lead, respectively, the African L3 branches in eastern Africa and the M and N L3 Eurasian branches in southeastern Asia. This model, that implies an earlier exit of modern humans out of Africa, has been contrasted with the results gathered independently by other disciplines.

## MATERIAL AND METHODS

### Sampling information

A total of 69 complete mtDNA genomes were sequenced in this study (Additional file 1: Table S1). They comprise the main African L haplogroups, excepting L6. To remedy this lack, 12 already published complete L6 sequences were included in our phylogenetic tree (Additional file 2: Figure S1). The different branches of haplogroup L3 are represented by 45 of these sequences. To establish the relative frequency of mtDNA macrohaplogroup L3 in the main African regions, a total of 25,203 partial and total mtDNA publicly available sequences were screened. Of them, 1,138 belong to our unpublished data (Additional file 1: Table S2). Similarly, to establish the relative frequency of Y-chromosome macrohaplogroup E in the main African regions, a total of 21,286 Y-chromosome publicly available African samples were screened. Of them, 737 belong to our unpublished data (Additional file 1: Table S2). All our samples were collected in the Canary Islands or Saudi Arabia from academic and health-care centers. The procedure of human population sampling adhered to the tenets of the Declaration of Helsinki and written consent was recorded from all participants before taking part in the study. The study underwent a formal review and was approved by the College of Medicine Ethical Committee of the King Saud University (proposal N° 09-659) and by the Ethics Committee for Human Research at the University of La Laguna (proposal NR157).

### MtDNA sequencing

Total DNA was isolated from buccal or blood samples using the POREGENE DNA isolation kit from Gentra Systems (Minneapolis, USA). PCR conditions and sequencing of mtDNA genome were as previously published [4]. Successfully amplified products were sequenced for both complementary strands using the DYEnamic™ETDye terminator kit (Amersham Biosciences). Samples run on MegaBACE™ 1000 (Amersham Biosciences) according to the manufacturer’s protocol. The 69 new complete mtDNA sequences have been deposited in GenBank under the accession numbers MF621062 to MF621130 (Additional file 1: Table S1).

### Previously published data compilation

Sequences belonging to specific mtDNA L haplogroups were obtained from public databases such as NCBI, MITOMAP, the-1000 Genomes Project and the literature. We searched for mtDNA lineages directly using diagnostic SNPs (http://www.mitomap.org/foswiki/bin/view/MITOMAP/WebHome), or by submitting short fragments including those diagnostic SNPs to a BLAST search (http://blast.st-va.ncbi.nlm.nih.gov/Blast.cgi). Haplotypes extracted from the literature were transformed into sequences using the HaploSearch program (http://www.haplosite.com/haplosearch/process/) [62]. Sequences were manually aligned and compared to the rCRS [63] with BioEdit Sequence Alignment program [64]. Haplogroup assignment was performed by hand, screening for diagnostic positions or motifs at both hypervariable and coding regions whenever possible.

Sequence alignment and haplogroup assignment were carried out twice by two independent researchers and any discrepancy resolved according to the PhyloTree database (Build 17; http://www.phylotree.org/) [65]. For the screening of the Y-chromosome haplogroup E, we considered samples as belonging to this haplogroup if in their analysis resulted positive for, at least, the diagnostic DE-YAP or E-M40, E-M96 markers.

### Phylogenetic analysis

The phylogenetic tree was constructed using the Network program, v4.6.1.2 using, in sequent order, the Reduced Median algorithm, Median Joining algorithm and Steiner (MP) algorithm [66].

Remaining reticulations were manually resolved. Haplogroup branches were named following the nomenclature proposed by the PhyloTree database [65]. Our coalescence ages were estimated by using statistics rho [67] and Sigma [68] and the calibration rate proposed by Soares et al.[6].

To calculate the total mean age of each haplogroup we recompiled all its different estimation ages from the literature without taken into account the mtDNA sequence segment analyzed, the mutation rate considered, or the inevitable partial overlapping of the samples used. In the cases where the same sample set was used to calculate its age by different methods, we always chose that performed with the rho statistic as the most generalized method (Additional file 1: Table S3). To calculate haplogroup mean coalescence ages for the non-recombining region of the Y-chromosome (NRY), we recompiled estimations preferably based on single nucleotide polymorphisms (SNPs) obtained by sequencing. When different mutation rates were used in the same study, we chose the age calculated with the slowest mutation rate (Additional file 1: Table S4).

### Phylogeographic analysis

In this study, we deal with the earliest periods of the out-of-Africa spread of modern humans and the likely return to Africa of the carriers of primitive mtDNA L3 and Y-chromosome E lineages. As the phylogeography of the different branches of these lineages has been exhaustively studied by other authors to assess more recent human movements on the continent, we focus here on its global distributions in the major African regions. For phylogeographic purposes, we divided the African continent into the following eight major regions: 1. Northwest Africa (including Morocco, West Sahara, Algeria and Tunisia), 2. Northeast Africa (including Libya and Egypt), 3. West Sahel (including Mauritania, Mali, and Niger), 4. East Sahel (including Chad, Sudan, Ethiopia, Somalia, and Eritrea), 5. West Guinea (including Senegambia, Guinea-Bissau, Guinea-Conakry, Sierra-Leona, Liberia, Ivory-Coast, Burkina-Faso, Ghana, Togo, Benin, and Nigeria), 6. Central Africa (including Cameroon, Central African Republic (CAR), Congo Democratic Republic (CDR), Congo-Brazzaville, Gabon and Equatorial Guinea), 7. East Guinea (including Uganda, Rwanda, Kenya, and Tanzania), 8. Southern Africa (including Angola, Zambia, Malawi, Mozambique, Zimbabwe, Botswana, Namibia and South African Republic (SAR)).

To evaluate the level of geographic structure of the mtDNA macrohaplogroup L3 and the Y-chromosome macrohaplogroup E in Africa, we performed AMOVA and K-means clustering analyses. We used the GenAlEx6.5 software to implement AMOVA and XLSTAT statistical software to perform the K-means clustering analysis. The possible association between the frequencies of mtDNA macrohaplogroup L3 and those of the Y-chromosome macrohaplogroup E, in the whole African continent, and in its principal geographic subdivisions, were tested by Pearson correlation analyses using the XLSTAT statistical software. As an important overlap exists among the expansion ages of the L3 branches with those of the African widespread mtDNA macrohaplogroup L2, the global frequencies of L2 have also been included in the majority of the phylogeographic analyses performed.

## RESULTS AND DISCUSSION

### Phylogeny and affinities of our African complete sequences

As a general rule, our 69 mtDNA complete sequences (Additional file 2: Figure S1) could be allocated into previously defined clades in the PhyloTree. Their closest affinities were with other sequences of the same haplogroups. Thus, our Kenyan L3a1a (Kn028) sequence shares tip mutations 514, 3796 and 4733 with a Tanzanian sequence (EF184630) but only 514 with a Somalian sequence (JN655813) of the same clade. The Sudanese L3b1a (Su238) sequence shares the very conservative transition at 12557 with an L3b sequence (KF055324) from an African-American glaucoma patient [69]. Our L3b1a2 (Su002) sequence has matches at 195, 12490 and 16311 with several African sequences (EU092669, EU092744, EU092795, EU092825, EU9355449) with which could make up a new branch, L3b1a2a, defined by these three transitions. In the same way, the L3f2a1 (Su004) has matches at mutated positions 6182, 8676, 9731, 12280, 12354 and 13105 with other published Senegalese sequences (JN655832, JN655841) with which could conform a new derived branch. It is expected that L sequences detected in the Canary Islands have their closest relatives in the African continent. Indeed, this is the case, for instance, for the L3d1b3 (Go764) sequence from La Gomera island that shares tip transitions 14040 and 16256 with an Ovimbundu isolate (KJ185837) from Angola [70]. However, unexpectedly, there are Canarian sequences as TF0005, allocated into the L3f1b subclade, that has its closest relatives in the Iberian Peninsula, sharing the 8994 transition with two Asturian L3f1b sequences (KJ959229, KJ959230) [71]. Similar is the case of the L3x2 (TF116) sequence from Tenerife that shares all its terminal variants(650, 7933, 8158, 15519, 16261) with sequences from Galicia (HQ675033, JN214446) and Andalusia (KT819228) instead of African sequences. Saudi Arabia has been identified as a universal receptor of mtDNA Eurasian lineages, which is also valid for the African female flow. Arab sequences belonging to the L3i1a (AR429) and L3x1a1 (AR260) haplogroups have their closest relatives with sequences JN655780 and DQ341067 respectively from the nearby Ethiopia, and the L3h1b1 (AR381) sequence is identical to an already published Yemeni isolate (KM986547). However, the L3h1b2 (AR221) sequence is most related to the JQ044990 lineage from Burkina Faso [72], with which shares particular transitions at 7424, 13194, 16192 and 16218 positions. The affinities of the Arab L1c2b1a’b (AR1252) with other sequences are the most striking.

This sequence, particularly characterized by the presence of an insertion of 11 nucleotides at the 16029 position in the control region, has an exact match with an L1 isolate from the Dominican Republic (DQ341059). Its closest relatives in Africa, although without that insertion, have been found in Angola (KJ185814) and Zambia (KJ185662) among Bantu-speakers [70]. The control region of this AR1252 isolate was previously published (KP960821). Concerning the less frequent L4, L5, and L6 clades, our L4b1a (Iv136) sequence from the Ivory Coast shares tip mutations 789, 7166 and 14935 with geographically close sequences (JQ044848, JQ045081) from Burkina Faso [72]. Likewise, the Arab L4a2 (AR1116) sequence is closely related to other African L4a2 sequences (EU092799, EU092800), and the L4b2a1 (AR197) isolate resulted identical to a sequence (KM986608) from Yemen [73]. From the analysis of partial sequences [53, 74], we can assure representatives of branches L4a1, L4a2 and L4b2 exist in Saudi Arabia. However, we have not yet detected sequences belonging to the large Sudanese L4b1b clade (Additional file 2: Figure S1). Published [53, 74] and unpublished data allow us to confirm that L5a is represented in Saudi Arabia by at least a lineage that has the following haplotype in, and nearby the coding region: 15884, 16093, 16129, 16148, 16166, 16187, 16189, 16223, 16265C, 16311, 16355, 16362/ 73, 152, 182, 195, 198, 247, 263, 315iC, 455i2T, 459iC, 513, 522dCA, 709, 750, 769, 825A, 851, 930. It represents 0.58% of the Saudi mtDNA gene pool. There is also a L5b lineage characterized by mutations at: 15927, 16111, 16129, 16148, 16166, 16187, 16189, 16223, 16233, 16254, 16265C, 16278, 16360, 16519/ 73, 195, 247, 249d, 263, 315iC, 459iC, 501, 535, 750, 769, and 825A, with minor presence (0.09%) in Saudi Arabia. For the same reason, although we lack complete L6 sequences, from partial sequencing and specific SNP analysis, we can confirm that L6a1 (0.13%) and L6b (0.09%) lineages are also present in the Saudi population [53, 74]. With 12 complete sequences, L3h is the best-represented haplogroup in our phylogeny (Additional file 2: Figure S1). In it, we have, provisionally, defined some new branches. We think that retromutation at 16223 position could define a Sudanese L3h1a1a branch. Two clades, L3h1a2a1a and L3h1a2a1b could be characterized by transitions 3892, 7705, 15346 and transitions 5108, 16165 respectively. An additional subclade, defined by transitions 7310, 13153, 14407 and transversion 9824A has been provisionally named as L3h1a2b1. Finally, after introducing the AR221 sequence, the old branch L3h1b2 would be characterized only by transitions 294, 8842, 9758, 12882, 13437, 16129, and 16362.

Our only discrepancy with the PhyloTree phylogeny, is about to the rare and old L5 clade. We have identified a new branch, provisionally named L5c (Additional file 2: Figure S1). With the information provided by this lineage, we think that the PhyloTree L5b node, that joints haplogroups L5b1 and L5b2 by sharing retromutations at 182, 13105 and 16311 positions and transition at 16254 position, lacks phylogenetic robustness. Instead, the PhyloTree L5b2 clade would be better considered a sister branch of our new Sudanese L5c sequence (Su412), both joined by the sharing of retromutation at the 195 position and transition at the 6527 and 11809 positions.

Lastly, it is worth mentioning that, despite the relatively small sample sizes employed, our coalescence age results fit well into the standard deviations calculated for the different haplogroups (Additional file 1: Table S3). Even so, mean age values for the L3 macrohaplogroup in Africa (71 ± 12 kya), that theoretically marks the lower bound time for the out-of-Africa of modern humans, falls short compared to those obtained from the fossil record in the Levant [75].

### The Eurasian origin of macrohaplogroup L3

The southern route hypothesis proposes that the Eurasian branches (M and N) of the macrohaplogroup L3 differentiated in or near the African continent and rapidly spread across the Asian peninsulas to reach Australia and Melanesia [45]. Under this assumption, it is expected that, in general, coalescence ages of haplogroups should decrease from Africa to Australia. However, we have demonstrated that this is not the case [53–55]. Just on the contrary, The oldest M and N haplogroups are detected in southern China and Australasia instead of India, and associations between longitudinal geographic distances and relative ages of M and N haplogroups run, against to expectation, westwards with younger haplogroup ages going to Africa [53, 54]. So, we confront a dilemma; it seems that two gravity centers of L3 expansion exist, one in Africa and the other in southeastern Asia. A geographic equidistant midpoint would situate the primitive radiation of L3 in India if a southern route were chosen by the African colonizers or above the Himalayas, between Tibet and Pamir, if the northern route was preferred. Furthermore, as the coalescence age of the African L3 branches and that of the Eurasian L3 (MN) are very similar (Table 1) and around 71 kya, the temporal and spatial midpoints might also coincide. As the group of modern humans that hypothetically turned back to Africa should include females and males, searching for the Y-chromosome phylogenetic and phylogeographic information might give us additional information. Indeed, an origin in Asia and return to Africa was proposed, long ago, for the Y-chromosome African haplogroup E [61]. This hypothesis was based on the derived state of its African YAP^+^ haplotypes 4 and 5 (haplogroup E) respect to the ancestral Asian YAP^+^ haplotype 3 (Haplogroup D). The later discovery of new markers evidenced that D and E were, in fact, sister branches of the YAP+ node. Haplogroup D showing the derived status for M174 and the ancestral status for M40 in Asia and, on the contrary, haplogroup E being characterized by M40 derived and M174 ancestral status all around Africa, so that the migratory sense between continents of both haplogroups could not be assured [76]. A few YAP^+^ individuals, ancestral for both markers, were detected in West Africa [77] and in Tibet [78]. Although assigned to the para-haplogroup DE*, its real ancestral state could not be confirmed. Likewise, a new mutation (P143) united the two other Eurasian haplogroups C and F as brothers and, in turn, DE and CF were united in an ancestral node defined by mutations M168 and M294 [79]. At first, the solution proposed to this complicated scenario was that two independent migrations out of Africa occurred, one marked by D and the other by the CF pair of lineages [80]. However, a new twist occurred after the discovery of more than 60,000 single nucleotide variants by next generation sequencing techniques. A most parsimonious interpretation of the Y-chromosome phylogeny constructed with these variants is that the predominant African haplogroup E arose outside the continent and back-migrated to Africa [59]. The DE split as a lower bound (69.0 ± 14.7 kya) and the radiation of E into Africa as an upper bound (65.5 ± 8.5 kya) are dates highly coincidental with those estimated for the mtDNA haplogroup L3 expansions (Table 1). Furthermore, the spatial distribution of the residual Y-chromosome haplogroup D in Asia is also a good indicator of the geographical location of the putative DE split. The highest frequency and diversity of D is in the Tibet region. Although it is also present, at low incidence, throughout southeast Asia, the other two centers with notable frequency are in Japan and the Andaman Islands [78, 81, 82], pointing to the existence of edge relic areas of which could be, long ago, a more wide distribution. There are not native D lineages in India, weakening the possibility that this subcontinent was the center of the DE partition and, therefore, taking the wind out of the southern route supporters. Most probably, the divide of the Y-chromosome D and E haplogroups occurred up to the Himalayas and in or westward to the Tibet which also coincides with the hypothetical bifurcation center proposed for the mtDNA L3 macrohaplogroup. As these coincidental female and male splits occurred during a glaciations time (70 - 100 Kya), it is reasonable to think that cold climatic conditions forced humans southwards and, confronted with the Himalayas, dispersed across southeastern and southwestern Asia. Most probably, this climatic change also obliged the Neanderthals to broaden its southern range and, therefore, to augment its geographic overlap with humans and, possibly, with Denisovans, outcompeting them in search of recourses (Figure 1). This southward retreat was stronger at the western side, as witnessed by the total occupation of the Levant by the Neanderthals around 70 kya [83] and the forced return of modern humans, carrying mtDNA L3 and Y-chromosome E basic lineages, to Africa. However, the tables turned around 20 ky later. Then were modern humans who advanced westwards from inner Asia displacing Neanderthals in its way, colonizing East Asia, South Asia, and Central Asia from where they reached the Levant around 50 kya [84] and Europe short after [85–87]. It is worth mentioning that this westward modern human colonization was also proposed from an archaeological perspective [87, 88]. Under this scenario, early modern humans had to leave Africa much earlier than the time frame proposed by the geneticists under the mitochondrial molecular clock restrictions [89]. In a conciliatory approach, we would fix this period at the L3’4 or even at the L3’4’6 mtDNA coalescence nodes. That is, around 80 to 100 kya (Table 1), in such a way that, at least, mutations 769, 1018 and 16311, that define the basic L3* lineage, occurred already out of Africa. In the same way, the exit of the companion men could be dated at the split of branch CDEF-M168 from B-M181 about 86-120 kya [59, 90]. However, given the inaccuracies of the molecular clock, we rather prefer to trust on the fossil and climatic records to establish the out of Africa of early modern humans across the Levant around 125 kya as the most favorable period.

**Table 1.**
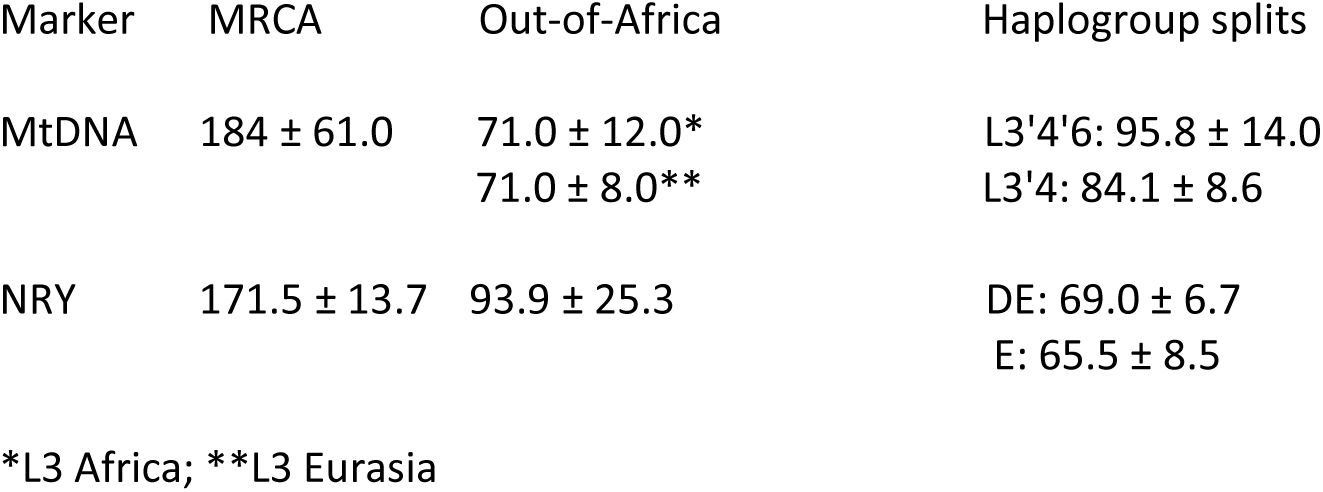
MtDNA and NRY mean values for MRCA, Out-of-Africa and haplogroup coalescences

| 1516 | Marker | MRCA | Out-of-Africa | Haplogroup splits |
| --- | --- | --- | --- | --- |
| 1517 |  |  |  |  |
| 1518 | MtDNA | 184 ± 61.0 | 71.0 ± 12.0* | L3'4'6: 95.8 ± 14.0 |
| 1519 |  |  | 71.0 ± 8.0** | L3'4: 84.1 ± 8.6 |
| 1520 |  |  |  |  |
| 1521 | NRY | 171.5 ± 13.7 | 93.9 ± 25.3 | DE: 69.0 ± 6.7 |
| 1522 |  |  |  | E: 65.5 ± 8.5 |
| 1523 |  |  |  |  |
| 1524 | *L3 Africa; **L3 Eurasia |  |  |  |

**Figure 1.**
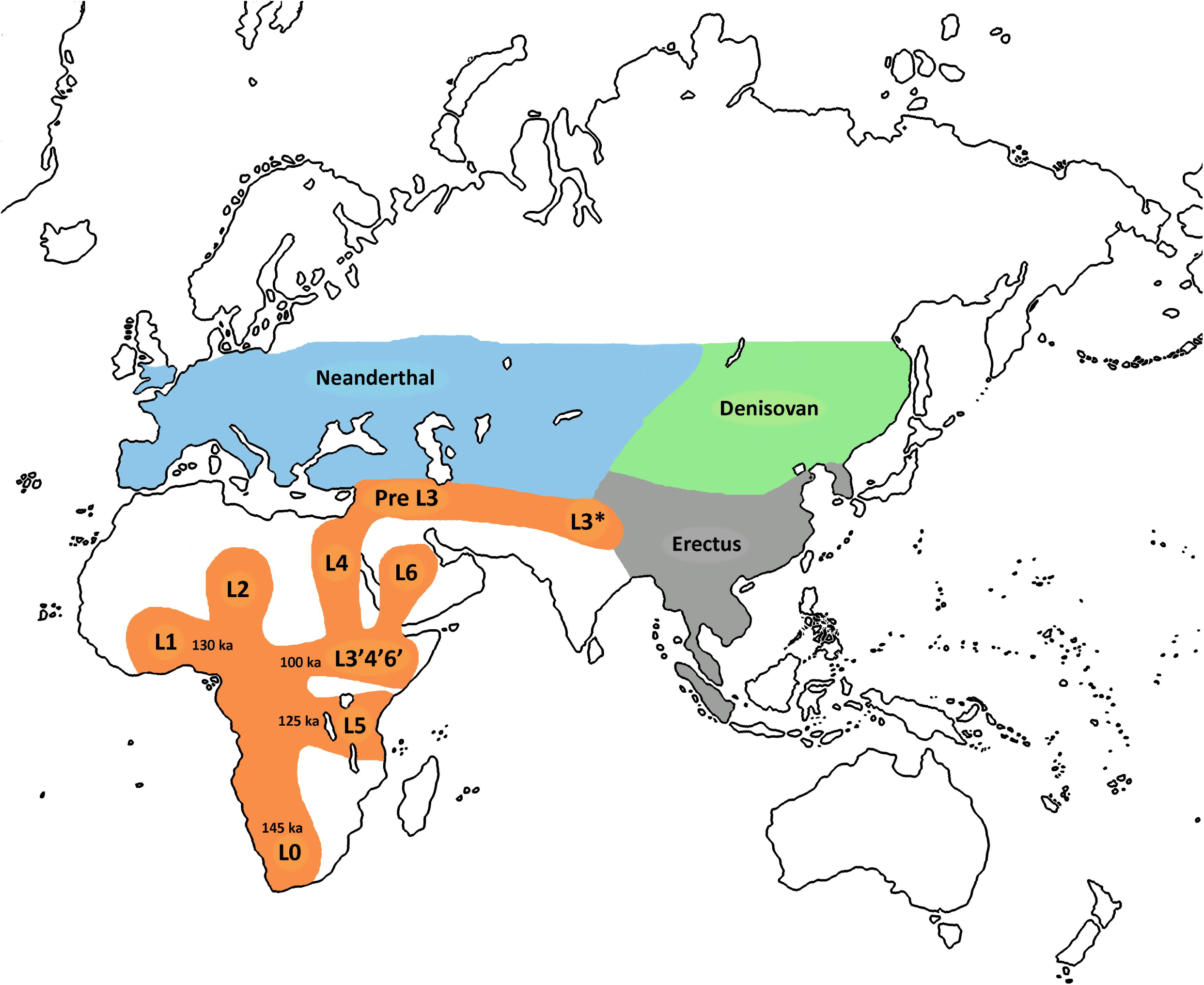
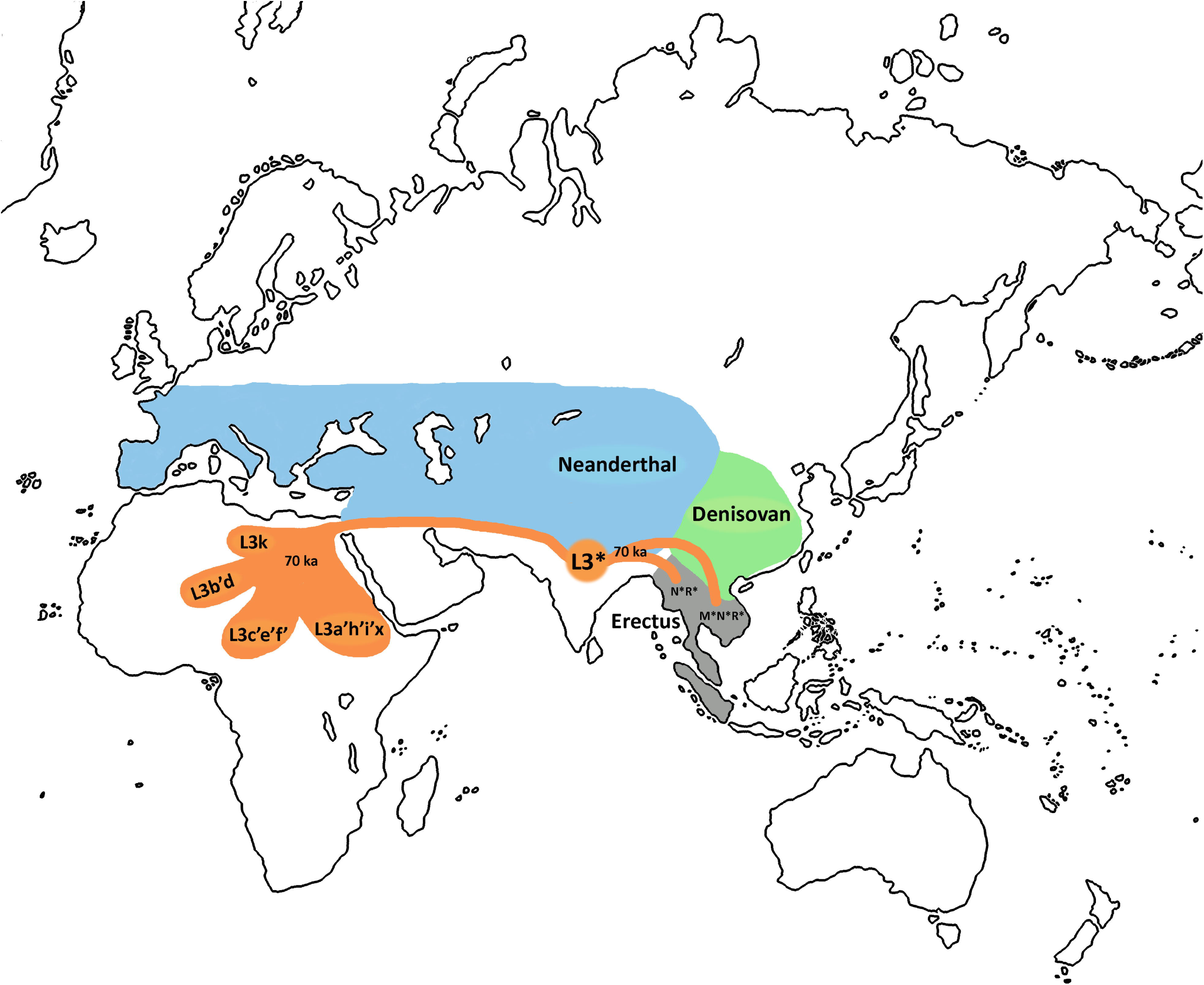
Geographic origin and dispersion of mtDNA L haplogroups: A. Sequential expansion of L haplogroups inside Africa and exit of the L3 precursor to Eurasia. B. Return to Africa and expansion to Asia of basic L3 lineages with subsequent differentiation in both continents.

### The phylogeography of the L3 and E lineages inside Africa

The global mean frequencies of the mtDNA and NRY haplogroups in the six main regions of the African continent are presented in Table 2. The mtDNA haplogroup L3 is more frequent in sub-Saharan Africa than in North Africa or the Sahel. In contrast, the Eurasian mtDNA haplogroups, including M1 and U6, are more frequent in northern Africa and the Sahel than in sub-Saharan Africa. In general, the Y-chromosome haplogroup E is more frequent in western than eastern Africa while the Eurasian Y haplogroups show a contrary trend. This geographic distribution confirms that the history of Africa is marked by multiple Eurasian migratory waves that pushed the first carriers of female haplogroup L3* and male haplogroup E* basic lineages inside Africa. It has been suggested that the few sub-Saharan haplogroups present in northern Africa are the result of recent historical incorporations [91]. Ancient DNA studies in the area seem to confirm this assumption both, in the northwest [92] and the northeast [93]. The fact that L3k, the only autochthonous L3 lineage in northern Africa, has only a residual presence in the area is also in favor of that suggestion [94]. Under this supposition, male E lineages, present nowadays in North Africa, would have reached the area as a secondary wave escorting Eurasian female lineages as M1 and U6. In fact, the main indigenous E clades in the region, E-M81 in the northwest, and E-M78 in the northeast are derived of haplogroup E-M35 which has also European Mediterranean and western Asian branches as E-V13 and E-V22 [95]. Against early studies that considered a Paleolithic implantation of E-M81 in the Maghreb [96, 97], it was suggested later that the low microsatellite diversity of this clade in northwest Africa could be better explained as the result of Neolithic or post-Neolithic gene flow episodes from the Near East [98]. However, after that, the discovery of a new sister branch of E-M81, named E-V257 [99], without Near Eastern roots but present in the European western and central Mediterranean shores and in Cameroon and Kenyan populations [99, 100], has weaken the suggested Levantine connection. Furthermore, E-M81 and E-M78 precursors are very old lineages that, respectively, accumulated 23 and 16 mutations in their basal branches. It has been reported that E-M78 radiated in eastern Africa in a time window between 20 and 15 kya but E-M81 did not, most probably because it was already in the Maghreb at that time. This would coincide with the expansion age in the area of the mtDNA U6a haplogroup [101]. Thus, a recent re-expansion after a large bottleneck would be the best explanation for the low variance of E-M81 in the present days [102]. The persistence of an even older male demographic substrate in this area has been evidenced by the detection in the region of representatives of the deepest Y phylogenetic clades A0 and A1a [103]. There is general consent in attributing an eastern African origin to the initial expansion of the NRY haplogroup E in the continent [100]. Curiously, ancestral E* lineages have been detected in the Arabian Peninsula [104] and the Levant [105]. Regardless of its origin, haplogroup E shows lower frequencies in northeastern Africa and the eastern Sahel compared with their counterparts in the West. Just the opposite occurs with the respective frequencies of Y Eurasian haplogroups in the same areas (Table 2), which points to later stronger Eurasian male gene flow throughout the northeastern side.

**Table 2.**
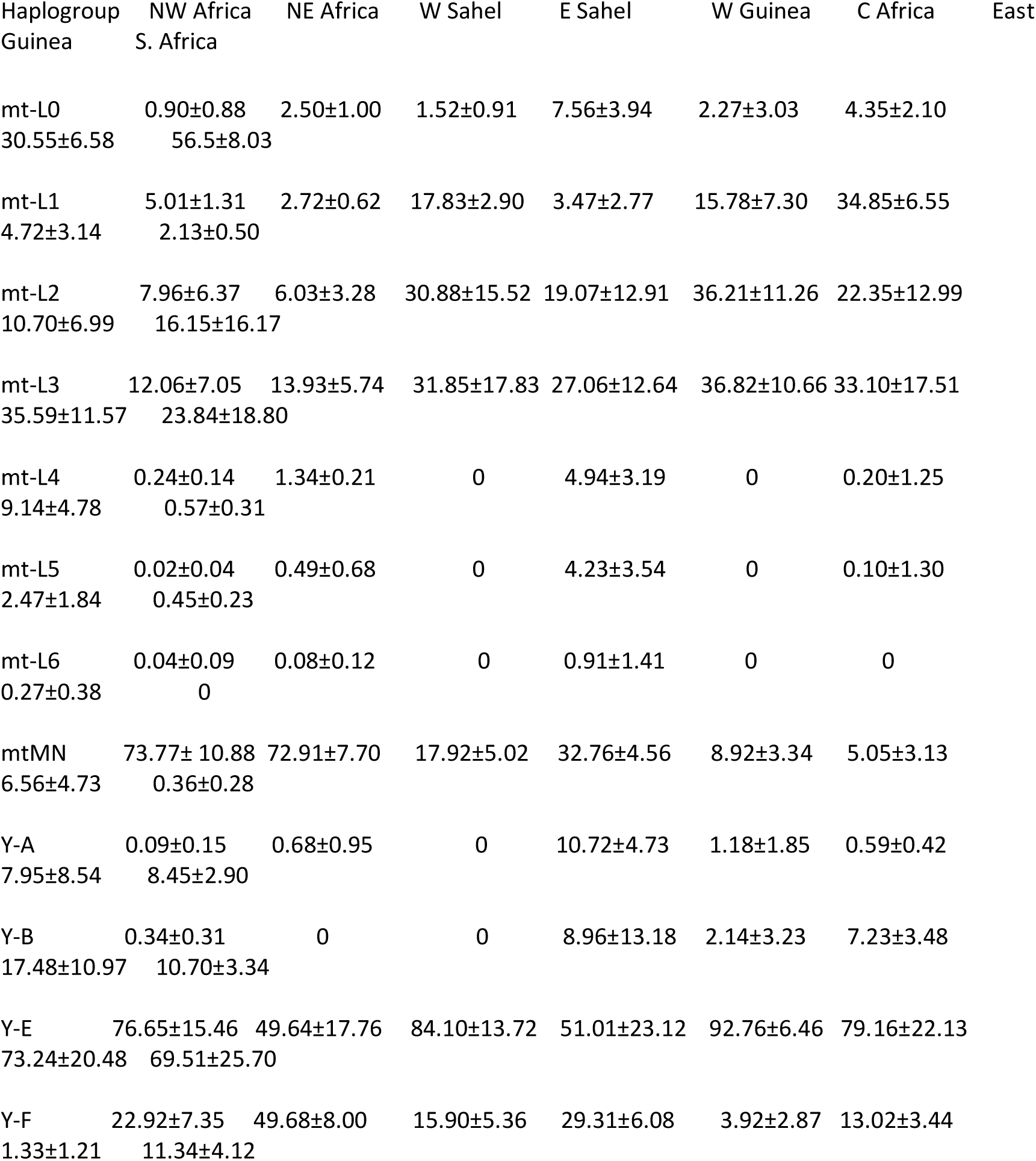
MtDNA and Y-Chromosome mean haplogroup frequencies in the major regions of the African continent

| 1538 | Haplogroup | NW Africa | NE Africa | W Sahel | E Sahel | W Guinea | C Africa | East |
| --- | --- | --- | --- | --- | --- | --- | --- | --- |
| 1539 | Guinea | S. Africa |  |  |  |  |  |  |
| 1540 |  |  |  |  |  |  |  |  |
| 1541 | mt-L0 | 0.90±0.88 | 2.50±1.00 | 1.52±0.91 | 7.56±3.94 | 2.27±3.03 | 4.35±2.10 |  |
| 1542 | 30.55±6.58 | 56.5±8.03 |  |  |  |  |  |  |
| 1543 |  |  |  |  |  |  |  |  |
| 1544 | mt-L1 | 5.01±1.31 | 2.72±0.62 | 17.83±2.90 | 3.47±2.77 | 15.78±7.30 | 34.85±6.55 |  |
| 1545 | 4.72±3.14 | 2.13±0.50 |  |  |  |  |  |  |
| 1546 |  |  |  |  |  |  |  |  |
| 1547 | mt-L2 | 7.96±6.37 | 6.03±3.28 | 30.88±15.52 | 19.07±12.91 | 36.21±11.26 | 22.35±12.99 |  |
| 1548 | 10.70±6.99 | 16.15±16.17 |  |  |  |  |  |  |
| 1549 |  |  |  |  |  |  |  |  |
| 1550 | mt-L3 | 12.06±7.05 | 13.93±5.74 | 31.85±17.83 | 27.06±12.64 | 36.82±10.66 | 33.10±17.51 |  |
| 1551 | 35.59±11.57 | 23.84±18.80 |  |  |  |  |  |  |

|  |  |  |  |  |  |  |  |
| --- | --- | --- | --- | --- | --- | --- | --- |
| 1552 |  |  |  |  |  |  |  |
| 1553 | mt-L4 | 0.24±0.14 | 1.34±0.21 | 0 | 4.94±3.19 | 0 | 0.20±1.25 |
| 1554 | 9.14±4.78 | 0.57±0.31 |  |  |  |  |  |
| 1555 |  |  |  |  |  |  |  |
| 1556 | mt-L5 | 0.02±0.04 | 0.49±0.68 | 0 | 4.23±3.54 | 0 | 0.10±1.30 |
| 1557 | 2.47±1.84 | 0.45±0.23 |  |  |  |  |  |
| 1558 |  |  |  |  |  |  |  |
| 1559 | mt-L6 | 0.04±0.09 | 0.08±0.12 | 0 | 0.91±1.41 | 0 | 0 |
| 1560 | 0.27±0.38 | 0 |  |  |  |  |  |
| 1561 |  |  |  |  |  |  |  |
| 1562 | mtMN | 73.77± 10.88 | 72.91±7.70 | 17.92±5.02 | 32.76±4.56 | 8.92±3.34 | 5.05±3.13 |
| 1563 | 6.56±4.73 | 0.36±0.28 |  |  |  |  |  |
| 1564 |  |  |  |  |  |  |  |
| 1565 | Y-A | 0.09±0.15 | 0.68±0.95 | 0 | 10.72±4.73 | 1.18±1.85 | 0.59±0.42 |
| 1566 | 7.95±8.54 | 8.45±2.90 |  |  |  |  |  |
| 1567 |  |  |  |  |  |  |  |
| 1568 | Y-B | 0.34±0.31 | 0 | 0 | 8.96±13.18 | 2.14±3.23 | 7.23±3.48 |
| 1569 | 17.48±10.97 | 10.70±3.34 |  |  |  |  |  |
| 1570 |  |  |  |  |  |  |  |
| 1571 | Y-E | 76.65±15.46 | 49.64±17.76 | 84.10±13.72 | 51.01±23.12 | 92.76±6.46 | 79.16±22.13 |
| 1572 | 73.24±20.48 | 69.51±25.70 |  |  |  |  |  |
| 1573 |  |  |  |  |  |  |  |
| 1574 | Y-F | 22.92±7.35 | 49.68±8.00 | 15.90±5.36 | 29.31±6.08 | 3.92±2.87 | 13.02±3.44 |
| 1575 | 1.33±1.21 | 11.34±4.12 |  |  |  |  |  |
| 1576 |  |  |  |  |  |  |  |
| 1577 |  |  |  |  |  |  |  |
| 1578 |  |  |  |  |  |  |  |
| 1579 |  |  |  |  |  |  |  |
| 1580 |  |  |  |  |  |  |  |
| 1581 |  |  |  |  |  |  |  |
| 1582 |  |  |  |  |  |  |  |

Detailed frequencies for mtDNA haplogroups L2 and L3 and Y-chromosome haplogroup E, all around the African continent, are listed in Additional file 1: Table S2. We have included L2 in the analysis because it was the sister clade of the composite eastern African node L3′4′6 that, through consecutive range expansions, promoted the exit of the L3 precursor out of Africa. Besides, inside Africa, several L2 derived spreads coincide in time with the later expansion of L3 branches in the continent. Furthermore, there is a suggestive positive association between the mean frequency estimates for L2 and the Y-chromosome haplogroup E across the major African regions (Table 2). In fact, there is a significant positive correlation between E haplogroup frequencies and both L3 (r = 0. 400; p < 0.0001) and L2 (r = 0.347; p < 0.0001) mtDNA haplogroup frequencies across Africa. Even more, this correlation is strongest when the E frequencies are compared with the sum of the two L2 and L3 frequencies (r = 0.477; p < 0.0001). However, the strength of this association varies in the different regions considered. Thus, the correlation is not significant at all in northern Africa, as could be expected from the picture commented above. It is barely significant in the Sahel region (r = 0.246; p = 0.045), but highly significant in the rest of the regions, with special intensity in southern Africa (r= 0.615; p < 0.0001). Nevertheless, these correlations are only slightly associated with geography as deduced from the AMOVA analysis that shows only a 4% of the variance due to differences among regions (Table 3). It seems that the E and L2/L3 expansions were strongest in western Sahel and western Guinea where they substituted the majority of the oldest mtDNA (L0 and L1) and Y-chromosome (A and B) lineages (Table 2). We applied the k-means clustering algorithm to our L3 and E frequency data (Additional file 1: Table S2). The consecutive partitioning of the samples into clusters has the objective of minimizing the variance within groups and augmenting the variance among them (Table 3). At k = 5, less than 20% of the variance is due to differences within clusters. At this level, the five classes obtained have centroid means for L3 and E that minimize the mean-square distance of the samples grouped to this center (Table 4 and Additional file 1: Tables S5 and S6). Class I, characterized by a relatively low frequency for both L3 and E haplogroups, joins the majority of the Khoesan-speaking groups from South Africa, Namibia, and Angola and the Hadza from Tanzania, but also several pygmy groups from Cameroon, Gabon, CAR and Congo as the Baka and the Babinga. In addition to their different geographic locations, these groups are also differentiated by the frequencies of other haplogroups. Thus, Khoesan-speaking samples harbor high frequencies of mtDNA L0d and L0k haplogroups and Y-chromosome A lineages, while the Central African Pygmies are characterized by the highest frequency of mtDNA L1c and Y-chromosome B-M60 lineages. In its turn, the Hadza share with pygmy groups the high percentages for B-M60 chromosomes and the highest frequency for mtDNA haplogroup L4 in the whole African continent. Other groups that belong to this class are the majority of Nilotes from Sudan and Uganda, characterized by their high frequencies of mtDNA haplogroup L2a1, and several Afro-Asiatic-speaking samples from Egypt and Sudan showing likewise high levels of L2a1 lineages but also of mtDNA L0a1 and Y-chromosome B representatives. Practically, the rest of the Khoesan-speaking samples fall into Class II characterized because, keeping a low frequency for L3 has an intermediate frequency for E (Table 4), pointing to a male-biased gene flow from western sub-Saharan Africans. At this class also belong the rest of the central Africa pygmies, including Sanga, Mbenzele, Biaka, and Mbuti, all still harboring high frequencies of Y B-M60 and L1c lineages. This class includes mainly northeast African and Ethiopian samples speaking Afro-Asiatic languages, some Nilo-Saharan speaking groups as the Fur from Sudan, the Anuak from Ethiopia or the Maasai from Kenya, and Bantu-speakers as the Shona from Zimbabwe and Botswana or the Tswana from Botswana and South Africa. However, the bulk of the northwestern African Afro-Asiatic speaking groups falls into Class III, defined by low frequencies of L3 and high frequencies of E (Table 4). A general low frequency for mtDNA haplogroup L2 is also a characteristic of this grouping. The majority of Berber and Tuareg samples belong to this class including the Gossi, Tamashek, and Douentza from Mali and the Gorom from Burkina Faso. Interestingly, the click-speaking Sandawe and the Nilo-Saharan speaking Datog from Tanzania are also assorted into Class III. From the maternal side, these Tanzanian samples are characterized by their relatively high frequencies of haplogroups L0a and L4. Nevertheless, the most abundant component of this class belongs to the Niger-Congo speakers including the majority of the Senegalese samples but also southern African specific Bantu-speakers as the Zulu and Xhosa. Curiously, the click-speaking Xeg Khoesan and Khwe belong to this cluster, pointing to substantial gene flow from Bantu-speaking immigrants. This fact was reported long ago for the Khwe that were found more closely linked to non-Khoesan-speaking, Bantu, populations [106]. Another class dominated by Niger-Congo speakers is Class IV, the largest of all. It groups samples that possess intermediate frequencies for L3 but high frequencies for E (Table 4). Linia and Kanembou from Chad, Rimaibe from Burkina-Faso, Songhai from Nigeria, and Masalit from Sudan are the only Nilo-Saharan speakers in this class. All the western African countries are represented by different Niger-Congo speaking groups, including the Bateke pygmies from Congo. There are also instances of eastern or southern African Bantu representatives. Finally, Class V groups those samples that present high frequencies for both, L3 and E, haplogroups (Table 4). Again, Niger-Congo speakers are the majority, although Nilo-Saharan from western countries as Menaka of Mali, Diffa of Niger and Kanuri from Nigeria likewise those of eastern countries as Bongor from Chad and Luo from Kenya are included in this class. The two best-represented countries are the western sub-Saharan Africa Nigeria and Cameroon which provided most of the Niger-Congo speaking samples. Nevertheless, there are also Bantu specific speakers from Kenya and southern Africa. This group includes the click-speakers Damara from Namibia and South Africa, who genetically have been associated to Bantu-speaking instead of to other Khoesan-speaking groups [107].

**Table 3.**
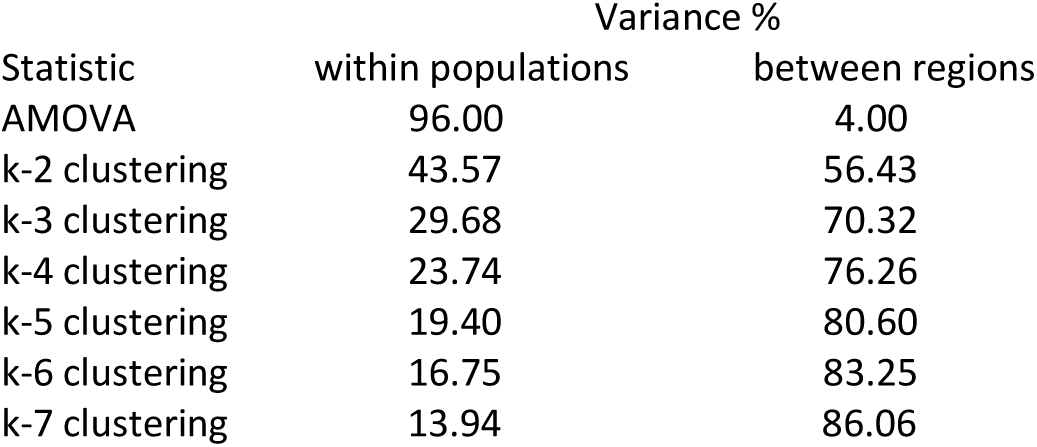
AMOVA and k-mean clustering results

**Table 4.** Frequency values for k-means centroids in 1 to 5 classes

| Clase | values L3/E | mt-L3 | Y-E |
| --- | --- | --- | --- |
| 1 | low/low | 16.9 | 25.9 |
| 2 | low/medium | 17.6 | 57.4 |
| 3 | low/high | 17.2 | 86.6 |
| 4 | medium/high | 36.2 | 92.9 |
| 5 | high/high | 55.0 | 87.2 |

The above-commented results show that the positive correlation found between Y-chromosome haplogroup E and mtDNA haplogroup L3 (and L2) lineages is neither strongly associated with the geography nor with language. It is better explained as the result of a gradual substitution of the most basal mtDNA (L0, L1, L5) and Y-chromosome (A, B) lineages by the phylogenetically younger clades L2 and L3, and E respectively throughout Africa. The data also point to important sex-biased dispersals between populations. These evident gene replacements in Africa have been mainly attributed to recent geographic range expansions of pastoralist and agriculturalist populations from eastern and western Africa at the expense of the hunter-gatherers inhabitants of the Central Africa rainforest [108–110], eastern African forested areas around the Great Lakes [111–114], and the semi-desert open spaces of South Africa [115–118]. Under our hypothesis of an early return to Africa from Eurasia of basic mtDNA L3 and Y-chromosome E lineages, and their expansion around 70 kya first into East and later into West Africa, those lineage replacements must have begun very early. It seems that in this first spread mtDNA haplogroup L2 was incorporated by favored female assimilation, whereas their hypothetical Y-chromosome haplogroup B counterparts were outcompeted by the incoming E chromosomes. An ancient expansion from a Central African source into eastern Africa at 70-50 kya has been associated with haplogroup L2 [119]. Likewise, an early expansion within Africa 60-80 kya involving L3 and, possibly, L2 was already detected long ago [120]. The last mentioned spread was considered the crucial event in the exit of modern humans from Africa into Eurasia. However, our proposition is that it signaled a backflow from Eurasia and subsequent expansion into Africa.

Finally, it seems interesting to point out that our hypothesis of an early return and subsequent expansion inside Africa of carriers of L3 and E haplogroups could help to explain, in a different way, the Neanderthal introgression detected in the western African Yoruba [121, 122], and in the northern African Tunisian Berbers [122].

### A new mtDNA model about the origin and dispersion of Homo sapiens

At mtDNA level, the sampling and data accumulated during the last thirty years, including those contributed by ancient DNA studies, allow us to propose a more detailed model of the origin and worldwide spread of modern humans than the ones proposed three decades ago. There are three fossil series in northwest, northeast, and southern Africa that chronologically and morphologically recapitulated the evolution of Homo sapiens from early archaic around 600 kya to early moderns by 200 kya [123]. The recent dating of Middle Stone Age tools (315 ± 34 kya) and early modern human fossils (286 ± 32 kya) from Jebel Irhoud in Morocco, places the emergence of our species, and of the Middle Stone Age, close in time and long before the age of about 200 kya previously suggested for the common origin of all humans in eastern Africa [124]. These data coincide in time with the existence of an old Y-chromosome lineage (A00) detected in samples of western-central African ascendance and dated 338 kya (95% CI: 237-581 kya), remarkably older than common estimates based on the Y-chromosome and mtDNA TMRCAs [125]. The fact that the following more divergent Y-chromosome A lineages (A0, A1a) also have a western-central African location, strongly supports this region as the origin of an ancestral human population from which the ancestors of early modern humans emerged [90, 103]. The most ancient splits and spreads of the mtDNA lineages also situated the hypothetical origin of all extant maternal lineages around this area. Although the earliest L0 clade diverged around 145 kya (Additional file 1: Table S3) and had its first expansions in southern Africa (L0d, L0k), the subsequent splits gave rise to L1 and L5 around 131 kya and 123 kya spreading to western and eastern Africa respectively. These long range African dispersions place its putative origin somewhere in Central Africa (Figure 1a). The same “centre-of-gravity” argument was used by other authors to suggest a Central African origin [126]. It is worth mentioning that while ancestral southern African Khoesan-speaking population still maintain high frequencies of primitive L0d and k lineages [94, 106, 127, 128], and that in the hunter-gatherer populations of central-western Africa the L1c haplogroup is dominant [108, 109], L5 in eastern Africa has today only a marginal presence [114, 129], most probably due to its displacement produced by more recent waves of better adapted incomers. The presence of L5 in southern Africa and eastern Mbuti pygmies [70, 109, 118, 127] is the result of later migrations. Most probably, next split, around 100 kya, also occurred in Central Africa resulting in sister clusters L2 and L3’4’6 that, respectively, produced initial westward and eastward expansions (Figure 1a). Although the oldest L2 lineages have been sampled in western Africa [130], today, as result of successive spreads inside the continent, this clade has a pan-African range [119]. In eastern Africa, the cluster L3’4’6 was the embryo of the full Eurasian maternal diversity. Its first split was haplogroup L6 that nowadays is a rare eastern lineage with a deep founder age (about 100 kya) but a rather recent expansion (about 25 kya). It has been found at frequencies below 1% in Egyptians [131], Somalis [132], Kenyan [133], and eastern Nilotes from Uganda [114]. Mean frequency rise in Ethiopia (3.15 ± 1.15 %) with a maximum (15.8%) in Ongota, an extinguishing linguistic isolate of uncertain adscription [129]. Outside Africa L6 has not been detected in the Levant [134]. It is present in the Arabian Peninsula at frequencies below 1% in Saudi samples but raises 12% in some Yemeni samples [135]. Attending to the L6 phylogeny (Additional file 2: Figure S1), it seems that not all the Yemeni lineages are a subset of the eastern African lineages as there is at least one for which its common node coincides with the expansion of the whole haplogroup. Based on its peculiar phylogeography, the possibility that L6 could have originated from the same out-of-Africa southern migration that colonized Eurasia was suggested [135]. If this were the case, this early L6 expansion would give genetic support to the reported presence of modern humans in the Arabian Peninsula, around 125 kya, based on archaeological evidence [12–14]. This suggestion also enjoys climatic support as this period coincides with humid environmental conditions in Arabia [136]. However, it seems that this possible human expansion did not extend beyond the Peninsula as L6 derived lineages have not yet been detected across Eurasia. The return to arid conditions, most probably, caused the decline of the populations carrying the L6 lineage that had to retreat to refuge areas as the highlands of Yemen and Ethiopia until more favorable conditions made possible their subsequent recovery in eastern Africa and Yemen. The long mutational stem that precedes the expansion of L6 (Additional file 2: Figure S1), would faithfully represent that strong and long bottleneck. Next phylogenetic bifurcation produced the ancestors of L3 and L4 haplogroups (Additional file 2: Figure S1). Nowadays, the highest frequencies and diversities of L4 are found in eastern Africa, but it has spread over the entire continent (Table 3). Besides, it has been detected at frequencies below 1% in the Levant [137], and the Arabian Peninsula [74, 138]. Most probably, as consequence of drift effects, some populations show outstanding frequencies of L4. In western Africa, Samoya (28.6%) and Kassena (21.2%) samples, speakers of the Gur linguistic family, stand out [72]. In Ethiopia, the cases of the Omotic-speaking Hamer (18.2%), the Cushitic-speaking Daasanach (22.2%), and the Nilotic-speaking Gumuz (24.0%) and Nyangatom (21.6%) are also remarkable [129, 139]. However, without any doubt, are the Tanzanian click-speaking Hadza (58%) and Sandawe (43%) whom show the highest values for L4 in Africa [111–113], this, together with the elevated frequencies that Hadza (50%) and Sandawe (15%) present for the Y-chromosome haplogroup B-M112 [140], points to human expansions from the North as those that most strongly influenced the gene pool of these groups. Attending to the age of bifurcation from L3 (around 95 kya), it could be thought that these L4 expansions occurred before our proposed return to Africa of L3 basic lineages. However, as the main spreads of its descendant clusters L4a (54.8 kya) and L4b (48.9 kya) [94, 138] had taken place around the same time window that the majority of the L3 and L2 branches in Africa, the most probable explanation is that improved climatic conditions after 60 kya motivated a global demographic growth on the African continent. Noticed that the evidence for an L3 first expansion in East Africa [89] is likewise in support of the out-of-Africa scenario than of a Eurasian back-flow as proposed here. We hypothetically situated the L3′4 node in northeast Africa or the Near East (Figure 1a) to allow an out-of-Africa of the pre-L3 clade. The Y-chromosome CDEF ancestor had to be its male counterpart. Other female and male lineages could have moved with them but, presumably gone extinct without contributing either to the maternal or paternal gene pools of the living human populations of Eurasia.

Under the scenario proposed here, early anatomically modern humans went out of Africa around 125 kya with a simple Middle Stone Age technology that was not superior to that manufactured by the Neanderthals. Favored by mild climatic conditions, these African pioneers progressed through West Asia and reached Central Asia overlapping in its way with the southern geographic range occupied by the Neanderthals. A new vision of the fossil and archaeological records of those regions [88, 141, 142] might uncover the path followed by those early African colonizers. At favorable conditions for both hominin groups, we might predict limited exchange of skills, lithic technology, and sex. However, when after 75 kya glacial environments became dominant, Neanderthals had to retreat southwards pushing out humans in its way. Confronted with the northern foothills of the Himalayas, humans moved in two directions, westwards to return to Africa, and eastwards to reach southeastern Asia across China (Figure 1b). The second part of this model has been already outlined in precedent articles [53–55].

## Additional files

Additional file 1: Table S1. Complete mtDNA macrohaplogroup L sequences. Table S2. Frequencies of mtDNA haplogroups L2 and L3 and Y-chromosome haplogroup E lineages across Africa. Table S3. Coalescence ages in thousand years (kya) with 95% coefficient intervals (CI), or standard deviations, for the main mitochondrial DNA African haplogroups. Table S4. Coalescence ages in thousand years (kya) with 95% coefficient intervals (CI), or standard deviations, for Y-chromosome most recent common ancestor (MRCA), the out-of-Africa event, and the splits of haplogroup DE and E. Table S5. k-means cluster results using African populations characterized by mtDNA L3 and Y-chromosome E haplogroup frequencies. Table S6. k-means cluster results using African populations characterized by mtDNA L2 and L3 and Y-chromosome E haplogroup frequencies.

Additional file 2: Figure S1. Phylogenetic tree of mtDNA macrohaplogroup L complete African sequences produced in this study.

## Acknowledgements

We are grateful to Dra. Ana M. González for her experimental contribution and bright ideas brought to this work. This research was supported by Grant n° CGL2010-16195 from the Spanish Ministerio de Ciencia e Innovación to JML.

## Availability of data and materials

The sequence set supporting the results of this article are available in the GenBank repository (MF621062 to MF621130), Additional file !: Table S1, and Additional file 2, Figure S1. References for the haplogroup frequencies used in this study are listed in Additional file 1, Table S2. All results obtained from our statistical analysis are presented in tables and figures of this article and in the additional files.

## Authors′ contributions

VMC conceived and designed the study, analyzed the data and wrote the manuscript. PM edited and submitted mtDNA sequences, designed figures, and contributed to the data analysis independently confirming the analysis results. KKAA carried out the sequencing of the Arabian and eastern African samples and made corrections on the manuscript. JML carried out the sequencing of the Canary Islands and western African samples, and contributed to the collection of published data and their analysis. All the authors read and approved the final manuscript.

## Authors′information

VMC is actually retired.

## Consent for publication

Not applicable.

## Ethics approval and consent to participate

The procedure of human population sampling adhered to the tenets of the Declaration of Helsinki. Written consent was recorded from all participants prior to taking part in the study. The study underwent formal review and was approved by the College of Medicine Ethical Committee of the King Saud University (proposal N°09-659), and by the Ethics Committee for Human Research at the University of La Laguna (proposal NR157).

## References

1 Cann RL, Stoneking M, Wilson AC: Mitochondrial DNA and human evolution. Nature, 325:31–6.

2 Stringer C: Why we are not all multiregionalists now. Trends in ecology \& evolution 2014, 29:248–251.

3 Ingman M, Kaessmann H, Pääbo S, Gyllensten U: Mitochondrial genome variation and the origin of modern humans. Nature 2000, 408:708–713.

4 Maca-Meyer N, González AM, Larruga JM, Flores C, Cabrera VM: Major genomic mitochondrial lineages delineate early human expansions. BMC genetics 2001, 2:1.

5 Herrnstadt C, Elson JL, Fahy E, Preston G, Turnbull DM, Anderson C, Ghosh SS, Olefsky JM, Beal MF, Davis RE, Howell N: Reduced-median-network analysis of complete mitochondrial DNA coding-region sequences for the major African, Asian, and European haplogroups. Am. J. Hum. Genet. 2002, 70:1152–71.

6 Soares P, Ermini L, Thomson N, Mormina M, Rito T, Röhl A, Salas A, Oppenheimer S, Macaulay V, Richards MB: Correcting for purifying selection: an improved human mitochondrial molecular clock. Am. J. Hum. Genet. 2009, 84:740–59.

7 Karmin M, Saag L, Vicente M, Sayres MAW, Järve M, Talas UG, Rootsi S, Ilumäe A-M, Mägi R, Mitt M, others: A recent bottleneck of Y chromosome diversity coincides with a global change in culture. Genome research 2015, 25:459–466.

8 Scally A, Durbin R: Revising the human mutation rate: implications for understanding human evolution. Nature Reviews Genetics 2012, 13:745–753.

9 Fu Q, Mittnik A, Johnson PL, Bos K, Lari M, Bollongino R, Sun C, Giemsch L, Schmitz R, Burger J, others: A revised timescale for human evolution based on ancient mitochondrial genomes. Current Biology 2013, 23:553–559.

10 Shea JJ, Bar-Yosef O: Who Were The Skhul/Qafzeh People? An Archaeological Perspective on Eurasia’s Oldest Modern Humans. WlpW ρκπ 2005:451–468.

11 Liu W, Martinón-Torres M, Cai Y, Xing S, Tong H, Pei S, Sier MJ, Wu X, Edwards RL, Cheng H, others: The earliest unequivocally modern humans in southern China. Nature 2015.

12 Armitage SJ, Jasim SA, Marks AE, Parker AG, Usik VI, Uerpmann H-P: The southern route “out of Africa”: evidence for an early expansion of modern humans into Arabia. Science 2011, 331:453–456.

13 Rose JI, Usik VI, Marks AE, Hilbert YH, Galletti CS, Parton A, Geiling JM, \vCern\y V, Morley MW, Roberts RG: The Nubian complex of Dhofar, Oman: an African middle stone age industry in southern Arabia. PLoS One 2011, 6:e28239.

14 Petraglia MD, Alsharekh A, Breeze P, Clarkson C, Crassard R, Drake NA, Groucutt HS, Jennings R, Parker AG, Parton A, others: Hominin dispersal into the Nefud desert and Middle Palaeolithic settlement along the Jubbah palaeolake, northern Arabia. PLoS One 2012, 7:e49840.

15 Sankararaman S, Patterson N, Li H, Pääbo S, Reich D: The date of interbreeding between Neandertals and modern humans. PLoS Genet 2012, 8:e1002947.

16 Kuhlwilm M, Gronau I, Hubisz MJ, de Filippo C, Prado-Martinez J, Kircher M, Fu Q, Burbano HA, Lalueza-Fox C, de La Rasilla M, others: Ancient gene flow from early modern humans into Eastern Neanderthals. Nature 2016, 530:429–433.

17 Xing J, Watkins WS, Hu Y, Huff CD, Sabo A, Muzny DM, Bamshad MJ, Gibbs RA, Jorde LB, Yu F: Genetic diversity in India and the inference of Eurasian population expansion. Genome Biol. 2010, 11:R113.

18 Pagani L, Lawson DJ, Jagoda E, Mörseburg A, Eriksson A, Mitt M, Clemente F, Hudjashov G, DeGiorgio M, Saag L, others: Genomic analyses inform on migration events during the peopling of Eurasia. Nature 2016, 538:238–242.

19 Quintana-Murci L, Semino O, Bandelt H-J, Passarino G, McElreavey K, Santachiara-Benerecetti AS: Genetic evidence of an early exit of Homo sapiens sapiens from Africa through eastern Africa. Nature genetics 1999, 23:437–441.

20 Quintana-Murci L, Chaix R, Wells RS, Behar DM, Sayar H, Scozzari R, Rengo C, Al-Zahery N, Semino O, Santachiara-Benerecetti AS, others: Where west meets east: the complex mtDNA landscape of the southwest and Central Asian corridor. The *American Journal of Human Genetics* 2004, 74:827–845.

21 Qin Z, Yang Y, Kang L, Yan S, Cho K, Cai X, Lu Y, Zheng H, Zhu D, Fei D, others: A mitochondrial revelation of early human migrations to the Tibetan Plateau before and after the last glacial maximum. American Journal of Physical Anthropology 2010, 143:555–569.

22 Kang L, Zheng H-X, Chen F, Yan S, Liu K, Qin Z, Liu L, Zhao Z, Li L, Wang X, others: mtDNA lineage expansions in Sherpa population suggest adaptive evolution

23 Wang C-C, Wang L-X, Shrestha R, Zhang M, Huang X-Y, Hu K, Jin L, Li H: Genetic structure of Qiangic populations residing in the western Sichuan corridor. PloS one 2014, 9:e103772.

24 Kong Q-P, Yao Y-G, Sun C, Bandelt H-J, Zhu C-L, Zhang Y-P: Phylogeny of east Asian mitochondrial DNA lineages inferred from complete sequences. Am. J. Hum. Genet. 2003, 73:671–6.

25 Tanaka M, Cabrera VM, González AM, Larruga JM, Takeyasu T, Fuku N, Guo L-J, Hirose R, Fujita Y, Kurata M, Shinoda K, Umetsu K, Yamada Y, Oshida Y, Sato Y, Hattori N, Mizuno Y, Arai Y, Hirose N, Ohta S, Ogawa O, Tanaka Y, Kawamori R, Shamoto-Nagai M, Maruyama W, Shimokata H, Suzuki R, Shimodaira H: Mitochondrial genome variation in eastern Asia and the peopling of Japan. Genome Res. 2004, 14:1832–50.

26 Derenko M, Malyarchuk B, Denisova G, Perkova M, Rogalla U, Grzybowski T, Khusnutdinova E, Dambueva I, Zakharov I: Complete mitochondrial DNA analysis of eastern Eurasian haplogroups rarely found in populations of northern Asia and eastern Europe. PloS one 2012, 7:e32179.

27 Metspalu M, Kivisild T, Metspalu E, Parik J, Hudjashov G, Kaldma K, Serk P, Karmin M, Behar DM, Gilbert MTP, others: Most of the extant mtDNA boundaries in south and southwest Asia were likely shaped during the initial settlement of Eurasia by anatomically modern humans. BMC genetics 2004, 5:1.

28 Gounder Palanichamy M, Sun C, Agrawal S, Bandelt H-J, Kong Q-P, Khan F, Wang C-Y, Chaudhuri TK, Palla V, Zhang Y-P: Phylogeny of mitochondrial DNA macrohaplogroup N in India, based on complete sequencing: implications for the peopling of South Asia. The American Journal of Human Genetics 2004, 75:966–978.

29 Cordaux R, Saha N, Bentley GR, Aunger R, Sirajuddin S, Stoneking M: Mitochondrial DNA analysis reveals diverse histories of tribal populations from India. European Journal of Human Genetics 2003, 11:253–264.

30 Chandrasekar A, Kumar S, Sreenath J, Sarkar BN, Urade BP, Mallick S, Bandopadhyay SS, Barua P, Barik SS, Basu D, others: Updating phylogeny of mitochondrial DNA macrohaplogroup m in India: dispersal of modern human in South Asian corridor. PloS one 2009, 4:e7447.

31 Hill C, Soares P, Mormina M, Macaulay V, Meehan W, Blackburn J, Clarke D, Raja JM, Ismail P, Bulbeck D, others: Phylogeography and ethnogenesis of aboriginal Southeast Asians. Molecular Biology and Evolution 2006, 23:2480–2491.

32 Summerer M, Horst J, Erhart G, Weißensteiner H, Schönherr S, Pacher D, Forer L, Horst D, Manhart A, Horst B, others: Large-scale mitochondrial DNA analysis in Southeast Asia reveals evolutionary effects of cultural isolation in the multi-ethnic population of Myanmar. BMC evolutionary biology 2014, 14:1.

33 Li Y-C, Wang H-W, Tian J-Y, Liu L-N, Yang L-Q, Zhu C-L, Wu S-F, Kong Q-P, Zhang Y-P: Ancient inland human dispersals from Myanmar into interior East Asia since the Late Pleistocene. Scientific reports 2015, 5.

34 Hill C, Soares P, Mormina M, Macaulay V, Clarke D, Blumbach PB, Vizuete-Forster M, Forster P, Bulbeck D, Oppenheimer S, others: A mitochondrial stratigraphy for island southeast Asia. The American Journal of Human Genetics 2007, 80:29–43.

35 Mona S, Grunz KE, Brauer S, Pakendorf B, Castr\’\i L, Sudoyo H, Marzuki S, Barnes RH, Schmidtke J, Stoneking M, others: Genetic admixture history of Eastern Indonesia as revealed by Y-chromosome and mitochondrial DNA analysis. Molecular Biology and Evolution 2009, 26:1865–1877.

36 Tabbada KA, Trejaut J, Loo J-H, Chen Y-M, Lin M, Mirazón-Lahr M, Kivisild T, de Ungria MCA: Philippine mitochondrial DNA diversity: a populated viaduct between Taiwan and Indonesia? Molecular biology and evolution 2010, 27:21–31.

37 Gunnarsdóttir ED, Li M, Bauchet M, Finstermeier K, Stoneking M: High-throughput sequencing of complete human mtDNA genomes from the Philippines. Genome research 2011, 21:1–11.

38 Tumonggor MK, Karafet TM, Hallmark B, Lansing JS, Sudoyo H, Hammer MF, Cox MP: The Indonesian archipelago: an ancient genetic highway linking Asia and the Pacific. Journal of human genetics 2013, 58:165–173.

39 Gomes SM, Bodner M, Souto L, Zimmermann B, Huber G, Strobl C, Röck AW, Achilli A, Olivieri A, Torroni A, others: Human settlement history between Sunda and Sahul: a focus on East Timor (Timor-Leste) and the Pleistocenic mtDNA diversity. BMC genomics 2015, 16:1.

40 Ingman M, Gyllensten U: Mitochondrial genome variation and evolutionary history of Australian and New Guinean aborigines. Genome Research 2003, 13: 1600–1606.

41 Van Holst Pellekaan SM, Ingman M, Roberts-Thomson J, Harding RM: Mitochondrial genomics identifies major haplogroups in Aboriginal Australians. American journal of physical anthropology 2006, 131:282–294.

42 Hudjashov G, Kivisild T, Underhill PA, Endicott P, Sanchez JJ, Lin AA, Shen P, Oefner P, Renfrew C, Villems R, others: Revealing the prehistoric settlement of Australia by Y chromosome and mtDNA analysis. Proceedings of the National Academy of Sciences 2007, 104:8726–8730.

43 Friedlaender JS, Friedlaender FR, Hodgson JA, Stoltz M, Koki G, Horvat G, Zhadanov S, Schurr TG, Merriwether DA: Melanesian mtDNA complexity. PLoS One 2007, 2:e248.

44 Duggan AT, Evans B, Friedlaender FR, Friedlaender JS, Koki G, Merriwether DA, Kayser M, Stoneking M: Maternal history of Oceania from complete mtDNA genomes: contrasting ancient diversity with recent homogenization due to the Austronesian expansion. The American Journal of Human Genetics 2014, 94:721–733.

45 Macaulay V, Hill C, Achilli A, Rengo C, Clarke D, Meehan W, Blackburn J, Semino O, Scozzari R, Cruciani F, others: Single, rapid coastal settlement of Asia revealed by analysis of complete mitochondrial genomes. Science 2005, 308:10341036.

46 Kong Q-P, Sun C, Wang H-W, Zhao M, Wang W-Z, Zhong L, Hao X-D, Pan H, Wang S-Y, Cheng Y-T, others: Large-scale mtDNA screening reveals a surprising matrilineal complexity in east Asia and its implications to the peopling of the region. Molecular biology and evolution 2011, 28:513–522.

47 Fernandes V, Alshamali F, Alves M, Costa MD, Pereira JB, Silva NM, Cherni L, Harich N, Cerny V, Soares P, others: The Arabian cradle: mitochondrial relicts of the first steps along the southern route out of Africa. The American Journal of Human Genetics 2012, 90:347–355.

48 Cordaux R, Weiss G, Saha N, Stoneking M: The northeast Indian passageway: a barrier or corridor for human migrations? Molecular Biology and Evolution 2004, 21:1525–1533.

49 Merriwether DA, Hodgson JA, Friedlaender FR, Allaby R, Cerchio S, Koki G, Friedlaender JS: Ancient mitochondrial M haplogroups identified in the Southwest Pacific. Proceedings of the National Academy of Sciences of the United States of America 2005, 102:13034–13039.

50 Van Holst Pellekaan S: Genetic evidence for the colonization of Australia. Quaternary International 2013, 285:44–56.

51 Pagani L, Schiffels S, Gurdasani D, Danecek P, Scally A, Chen Y, Xue Y, Haber M, Ekong R, Oljira T, others: Tracing the route of modern humans out of Africa by using 225 human genome sequences from Ethiopians and Egyptians. The American Journal of Human Genetics 2015, 96:986–991.

52 Sankararaman S, Mallick S, Dannemann M, Prüfer K, Kelso J, Pääbo S, Patterson N, Reich D: The genomic landscape of Neanderthal ancestry in present-day humans. Nature 2014, 507:354–357.

53 Fregel R, Cabrera V, Larruga JM, Abu-Amero KK, González AM: Carriers of Mitochondrial DNA Macrohaplogroup N Lineages Reached Australia around 50.0 Years Ago following a Northern Asian Route. PloS one 2015, 10:e0129839.

54 Marrero P, Abu-Amero KK, Larruga JM, Cabrera VM: Carriers of human mitochondrial DNA macrohaplogroup M colonized India from southeastern Asia. bioRxiv 2016:047456.

55 Larruga JM, Marrero P, Abu-Amero KK, Golubenko MV, Cabrera VM: Carriers of mitochondrial DNA macrohaplogroup R colonized Eurasia and Australasia from a southeast Asia core area. BMC evolutionary biology 2017, 17:115.

56 Magoon GR, Banks RH, Rottensteiner C, Schrack BE, Tilroe VO, Grierson AJ: Generation of high-resolution a priori Y-chromosome phylogenies using “next-generation” sequencing data. bioRxiv 2013:000802.

57 Karafet TM, Mendez FL, Sudoyo H, Lansing JS, Hammer MF: Improved phylogenetic resolution and rapid diversification of Y-chromosome haplogroup K-M526 in Southeast Asia. European Journal of Human Genetics 2014.

58 Hallast P, Batini C, Zadik D, Delser PM, Wetton JH, Arroyo-Pardo E, Cavalleri GL, de Knijff P, Bisol GD, Dupuy BM, others: The Y-chromosome tree bursts into leaf: 13.0 high-confidence SNPs covering the majority of known clades. Molecular biology and evolution 2015, 32:661–673.

59 Poznik GD, Xue Y, Mendez FL, Willems TF, Massaia A, Sayres MAW, Ayub Q, McCarthy SA, Narechania A, Kashin S, others: Punctuated bursts in human male demography inferred from 1,244 worldwide Y-chromosome sequences. Nature genetics 2016, 48:593–599.

60 Groucutt HS, Petraglia MD, Bailey G, Scerri EM, Parton A, Clark-Balzan L, Jennings RP, Lewis L, Blinkhorn J, Drake NA, others: Rethinking the dispersal of Homo sapiens out of Africa. Evolutionary Anthropology: Issues, News, and Reviews 2015, 24:149–164.

61 Hammer MF, Karafet T, Rasanayagam A, Wood ET, Altheide TK, Jenkins T, Griffiths RC, Templeton AR, Zegura SL: Out of Africa and back again: nested cladistic analysis of human Y chromosome variation. Mol. Biol. Evol. 1998, 15:427–41.

62 Fregel R, Delgado S: HaploSearch: a tool for haplotype-sequence two-way transformation. Mitochondrion 2011, 11:366–367.

63 Andrews RM, Kubacka I, Chinnery PF, Lightowlers RN, Turnbull DM, Howell N: Reanalysis and revision of the Cambridge reference sequence for human mitochondrial DNA. Nature genetics 1999, 23:147–147.

64 Hall TA: BioEdit: a user-friendly biological sequence alignment editor and analysis program for Windows 95/98/NT. In Nucleic acids symposium series. 1999, 41:95–98.

65 Van Oven M, Kayser M: Updated comprehensive phylogenetic tree of global human mitochondrial DNA variation. Hum. Mutat. 2009, 30:E386–94.

66 Bandelt H-J, Forster P, Röhl A: Median-joining networks for inferring intraspecific phylogenies. Molecular biology and evolution 1999, 16:37–48.

67 Forster P, Harding R, Torroni A, Bandelt HJ: Origin and evolution of Native American mtDNA variation: a reappraisal. Am. J. Hum. Genet. 1996, 59:935–45.

68 Saillard J, Forster P, Lynnerup N, Bandelt HJ, Nørby S: mtDNA variation among Greenland Eskimos: the edge of the Beringian expansion. Am. J. Hum. Genet. 2000, 67:718–26.

69 Collins DW, Gudiseva HV, Trachtman BT, Jerrehian M, Gorry T, Merritt WT, Rhodes AL, Sankar PS, Regina M, Miller-Ellis E, O’Brien JM: Mitochondrial sequence variation in African-American primary open-angle glaucoma patients. PLoS ONE 2013, 8:e76627.

70 Barbieri C, Vicente M, Oliveira S, Bostoen K, Rocha J, Stoneking M, Pakendorf B: Migration and interaction in a contact zone: mtDNA variation among Bantu-speakers in southern Africa. PloS one 2014, 9:e99117.

71 Pardinas AF, Mart’\inez JL, Roca A, Garc’\ia-Vazquez E, López B: Over the sands and far away: interpreting an Iberian mitochondrial lineage with ancient Western African origins. American Journal of Human Biology 2014, 26:777–783.

72 Barbieri C, Whitten M, Beyer K, Schreiber H, Li M, Pakendorf B: Contrasting maternal and paternal histories in the linguistic context of Burkina Faso. Molecular biology and evolution 2011, 29:1213–1223.

73 Vyas DN, Kitchen A, Miró-Herrans AT, Pearson LN, Al-Meeri A, Mulligan CJ: Bayesian analyses of Yemeni mitochondrial genomes suggest multiple migration events with Africa and Western Eurasia. American journal of physical anthropology 2015.

74 Abu-Amero KK, Larruga JM, Cabrera VM, González AM: Mitochondrial DNA structure in the Arabian Peninsula. BMC evolutionary biology 2008, 8:1.

75 Mellars P, Gori KC, Carr M, Soares PA, Richards MB: Genetic and archaeological perspectives on the initial modern human colonization of southern Asia. Proc. Natl. Acad. Sci. U.S.A. 2013, 110:10699–704.

76 Underhill PA, Passarino G, Lin AA, Shen P, Lahr MM, Foley RA, Oefner PJ, Cavalli-Sforza LL: The phylogeography of Y chromosome binary haplotypes and the origins of modern human populations. Annals of human genetics 2001, 65:43–62.

77 Weale ME, Shah T, Jones AL, Greenhalgh J, Wilson JF, Nymadawa P, Zeitlin D, Connell BA, Bradman N, Thomas MG: Rare deep-rooting Y chromosome lineages in humans: lessons for phylogeography. Genetics 2003, 165:229–234.

78 Shi H, Zhong H, Peng Y, Dong Y-L, Qi X-B, Zhang F, Liu L-F, Tan S-J, Ma RZ, Xiao C-J, others: Y chromosome evidence of earliest modern human settlement in East Asia and multiple origins of Tibetan and Japanese populations. BMC biology 2008, 6:1.

79 Karafet TM, Mendez FL, Meilerman MB, Underhill PA, Zegura SL, Hammer MF: New binary polymorphisms reshape and increase resolution of the human Y chromosomal haplogroup tree. Genome Res. 2008, 18:830–8.

80 Underhill P a, Kivisild T: Use of y chromosome and mitochondrial DNA population structure in tracing human migrations. Annual review of genetics 2007, 41:539–64.

81 Su B, Xiao C, Deka R, Seielstad MT, Kangwanpong D, Xiao J, Lu D, Underhill P, Cavalli-Sforza L, Chakraborty R, others: Y chromosome haplotypes reveal prehistorical migrations to the Himalayas. Human genetics 2000, 107:582–590.

82 Thangaraj K, Singh L, Reddy AG, Rao VR, Sehgal SC, Underhill PA, Pierson M, Frame IG, Hagelberg E: Genetic affinities of the Andaman Islanders, a vanishing human population. Curr. Biol. 2003, 13:86–93.

83 Shea JJ: Transitions or turnovers? Climatically-forced extinctions of Homo sapiens and Neanderthals in the east Mediterranean Levant. Quaternary Science Reviews 2008, 27:2253–2270.

84 Hershkovitz I, Marder O, Ayalon A, Bar-Matthews M, Yasur G, Boaretto E, Caracuta V, Alex B, Frumkin A, Goder-Goldberger M, others: Levantine cranium from Manot Cave (Israel) foreshadows the first European modern humans. Nature 2015.

85 Rougier H, Milota S, Rodrigo R, Gherase M, Sarcina L, Moldovan O, Zilhão J, Constantin S, Franciscus RG, Zollikofer CPE, Ponce de León M, Trinkaus E: Peçtera cu Oase 2 and the cranial morphology of early modern Europeans. Proc. Natl. Acad. Sci. U.S.A. 2007, 104:1165–70.

86 Benazzi S, Slon V, Talamo S, Negrino F, Peresani M, Bailey SE, Sawyer S, Panetta D, Vicino G, Starnini E, others: The makers of the Protoaurignacian and implications for Neandertal extinction. Science 2015, 348:793–796.

87 Marks AE: Comments after four decades of research on the Middle to Upper Paleolithic transition. Mitteilungen der Gesellschaft für Urgeschichte 2005, 14:81–86.

88 Otte M: Central Asia as a Core Area: Iran as an Origin for the.

89 Soares P, Alshamali F, Pereira JB, Fernandes V, Silva NM, Afonso C, Costa MD, Musilová E, Macaulay V, Richards MB, Cerny V, Pereira L: The Expansion of mtDNA Haplogroup L3 within and out of Africa. Mol. Biol. Evol. 2012, 29:915–27.

90 Scozzari R, Massaia A, Trombetta B, Bellusci G, Myres NM, Novelletto A, Cruciani F: An unbiased resource of novel SNP markers provides a new chronology for the human Y chromosome and reveals a deep phylogenetic structure in Africa. Genome research 2014, 24:535–544.

91 Harich N, Costa MD, Fernandes V, Kandil M, Pereira JB, Silva NM, Pereira L: The trans-Saharan slave trade-clues from interpolation analyses and high-resolution characterization of mitochondrial DNA lineages. BMC evolutionary biology 2010, 10:138.

92 Kefi R, Hechmi M, Naouali C, Jmel H, Hsouna S, Bouzaid E, Abdelhak S, Beraud-Colomb E, Stevanovitch A: On the origin of Iberomaurusians: new data based on ancient mitochondrial DNA and phylogenetic analysis of Afalou and Taforalt populations. Mitochondrial DNA Part A 2016:1–11.

93 Schuenemann VJ, Peltzer A, Welte B, Van Pelt WP, Molak M, Wang C-C, Furtwängler A, Urban C, Reiter E, Nieselt K, others: Ancient Egyptian mummy genomes suggest an increase of Sub-Saharan African ancestry in post-Roman periods. Nature communications 2017, 8.

94 Behar DM, Villems R, Soodyall H, Blue-Smith J, Pereira L, Metspalu E, Scozzari R, Makkan H, Tzur S, Comas D, others: The dawn of human matrilineal diversity. The American Journal of Human Genetics 2008, 82:1130–1140.

95 Cruciani F, La Fratta R, Trombetta B, Santolamazza P, Sellitto D, Colomb EB, Dugoujon J-M, Crivellaro F, Benincasa T, Pascone R, others: Tracing past human male movements in northern/eastern Africa and western Eurasia: new clues from Y-chromosomal haplogroups E-M78 and J-M12. Molecular biology and evolution 2007, 24:1300–1311.

96 Bosch E, Calafell F, Comas D, Oefner PJ, Underhill PA, Bertranpetit J: Highresolution analysis of human Y-chromosome variation shows a sharp discontinuity and limited gene flow between northwestern Africa and the Iberian Peninsula. The *American Journal of Human Genetics* 2001, 68:1019–1029.

97 Flores C, Maca-Meyer N, Perez JA, Hernandez M, Cabrera VM: Y-chromosome differentiation in Northwest Africa. Human biology 2001, 73:513–524.

98 Arredi B, Poloni ES, Paracchini S, Zerjal T, Fathallah DM, Makrelouf M, Pascali VL, Novelletto A, Tyler-Smith C: A predominantly neolithic origin for Y-chromosomal DNA variation in North Africa. Am. J. Hum. Genet. 2004, 75:338–45.

99 Trombetta B, Cruciani F, Sellitto D, Scozzari R: A new topology of the human Y chromosome haplogroup E1b1 (E-P2) revealed through the use of newly characterized binary polymorphisms. PLoS One 2011, 6:e16073.

100 Trombetta B, D’Atanasio E, Massaia A, Ippoliti M, Coppa A, Candilio F, Coia V, Russo G, Dugoujon J-M, Moral P, others: Phylogeographic refinement and large scale genotyping of human Y chromosome haplogroup E provide new insights into the dispersal of early pastoralists in the African continent. Genome biology and evolution 2015, 7:1940–1950.

101 Secher B, Fregel R, Larruga JM, Cabrera VM, Endicott P, Pestano JJ, González AM: The history of the North African mitochondrial DNA haplogroup U6 gene flow into the African, Eurasian and American continents. BMC Evol. Biol. 2014, 14: 109.

102 Semino O, Magri C, Benuzzi G, Lin AA, Al-Zahery N, Battaglia V, Maccioni L, Triantaphyllidis C, Shen P, Oefner PJ, others: Origin, diffusion, and differentiation of Y-chromosome haplogroups E and J: inferences on the neolithization of Europe and later migratory events in the Mediterranean area. The American Journal of Human Genetics 2004, 74:1023–1034.

103 Cruciani F, Trombetta B, Massaia A, Destro-Bisol G, Sellitto D, Scozzari R: A revised root for the human Y chromosomal phylogenetic tree: the origin of patrilineal diversity in Africa. The American Journal of Human Genetics 2011, 88:814–818.

104 Abu-Amero KK, Hellani A, González AM, Larruga JM, Cabrera VM, Underhill PA: Saudi Arabian Y-Chromosome diversity and its relationship with nearby regions. BMC genetics 2009, 10:1.

105 Zalloua PA, Xue Y, Khalife J, Makhoul N, Debiane L, Platt DE, Royyuru AK, Herrera RJ, Hernanz DFS, Blue-Smith J, others: Y-chromosomal diversity in Lebanon is structured by recent historical events. The American Journal of Human Genetics 2008, 82:873–882.

106 Chen Y-S, Olckers A, Schurr TG, Kogelnik AM, Huoponen K, Wallace DC: mtDNA variation in the South African Kung and Khwe—and their genetic relationships to other African populations. The American Journal of Human Genetics 2000, 66:1362–1383.

107 Soodyall H, Makkan H, Haycock P, Naidoo T: The genetic prehistory of the Khoe and San. Southern African Humanities 2008, 20:37–48.

108 Quintana-Murci L, Quach H, Harmant C, Luca F, Massonnet B, Patin E, Sica L, Mouguiama-Daouda P, Comas D, Tzur S, others: Maternal traces of deep common ancestry and asymmetric gene flow between Pygmy hunter-gatherers and Bantu-speaking farmers. Proceedings of the National Academy of Sciences 2008, 105:15961601.

109 Batini C, Lopes J, Behar DM, Calafell F, Jorde LB, Van der Veen L, Quintana-Murci L, Spedini G, Destro-Bisol G, Comas D: Insights into the demographic history of African Pygmies from complete mitochondrial genomes. Molecular Biology and Evolution 2010, 28:1099–1110.

110 Anagnostou P, Battaggia C, Capocasa M, Boschi I, Brisighelli F, Batini C, Spedini G, Destro-Bisol G: Reevaluating a Model of Gender-Biased Gene Flow among Sub-Saharan Hunter-Gatherers and Farmers. Human biology 2013, 85:597–606.

111 Knight A, Underhill PA, Mortensen HM, Zhivotovsky LA, Lin AA, Henn BM, Louis D, Ruhlen M, Mountain JL: African Y chromosome and mtDNA divergence provides insight into the history of click languages. Current Biology 2003, 13:464473.

112 Henn BM, Gignoux C, Lin AA, Oefner PJ, Shen P, Scozzari R, Cruciani F, Tishkoff SA, Mountain JL, Underhill PA: Y-chromosomal evidence of a pastoralist migration through Tanzania to southern Africa. Proceedings of the National Academy of Sciences 2008, 105:10693–10698.

113 Tishkoff SA, Gonder MK, Henn BM, Mortensen H, Knight A, Gignoux C, Fernandopulle N, Lema G, Nyambo TB, Ramakrishnan U, others: History of click-speaking populations of Africa inferred from mtDNA and Y chromosome genetic variation. Molecular biology and evolution 2007, 24:2180–2195.

114 Gomes V, Pala M, Salas A, Álvarez-Iglesias V, Amorim A, Gómez-Carballa A, Carracedo Á, Clarke DJ, Hill C, Mormina M, others: Mosaic maternal ancestry in the Great Lakes region of East Africa. Human genetics 2015, 134:1013.

115 Salas A, Richards M, de La Fe T, Lareu M-V, Sobrino B, Sánchez-Diz P, Macaulay V, Carracedo Á: The making of the African mtDNA landscape. The American Journal of Human Genetics 2002, 71:1082–1111.

116 Coelho M, Sequeira F, Luiselli D, Beleza S, Rocha J: On the edge of Bantu expansions: mtDNA, Y chromosome and lactase persistence genetic variation in southwestern Angola. BMC evolutionary biology 2009, 9:80.

117 Pour NA, Plaster CA, Bradman N: Evidence from Y-chromosome analysis for a late exclusively eastern expansion of the Bantu-speaking people. European Journal of Human Genetics 2013, 21:423–429.

118 Marks SJ, Montinaro F, Levy H, Brisighelli F, Ferri G, Bertoncini S, Batini C, Busby GB, Arthur C, Mitchell P, others: Static and moving frontiers: the genetic landscape of Southern African Bantu-speaking populations. Molecular biology and evolution 2014, 32:29–43.

119 Silva M, Alshamali F, Silva P, Carrilho C, Mandlate F, Trovoada MJ, \vCern\’y V, Pereira L, Soares P: 60,000 years of interactions between Central and Eastern Africa documented by major African mitochondrial haplogroup L2. Scientific reports 2015, 5.

120 Watson E, Forster P, Richards M, Bandelt HJ: Mitochondrial footprints of human expansions in Africa. Am. J. Hum. Genet. 1997, 61:691–704.

121 Prüfer K, Racimo F, Patterson N, Jay F, Sankararaman S, Sawyer S, Heinze A, Renaud G, Sudmant PH, de Filippo C, others: The complete genome sequence of a Neanderthal from the Altai Mountains. Nature 2014, 505:43–49.

122 Sánchez-Quinto F, Botigué LR, Civit S, Arenas C, Ávila-Arcos MC, Bustamante CD, Comas D, Lalueza-Fox C: North African populations carry the signature of admixture with Neandertals. PLoS One 2012, 7:e47765.

123 Bräuer G: The origin of modern anatomy: by speciation or intraspecific evolution? Evolutionary Anthropology: Issues, News, and Reviews 2008, 17:22–37.

124 Richter D, Grün R, Joannes-Boyau R, Steele TE, Amani F, Rué M, Fernandes P, Raynal J-P, Geraads D, Ben-Ncer A, others: The age of the hominin fossils from Jebel Irhoud, Morocco, and the origins of the Middle Stone Age. Nature 2017, 546:293296.

125 Mendez FL, Krahn T, Schrack B, Krahn A-M, Veeramah KR, Woerner AE, Fomine FLM, Bradman N, Thomas MG, Karafet TM, others: An African American paternal lineage adds an extremely ancient root to the human Y chromosome phylogenetic tree. The American Journal of Human Genetics 2013, 92:454–459.

126 Rito T, Richards MB, Fernandes V, Alshamali F, Cerny V, Pereira L, Soares P: The first modern human dispersals across Africa. PLoS ONE 2013, 8:e80031.

127 Barbieri C, Güldemann T, Naumann C, Gerlach L, Berthold F, Nakagawa H, Mpoloka SW, Stoneking M, Pakendorf B: Unraveling the complex maternal history of Southern African Khoisan populations. American journal of physical anthropology 2014, 153:435–448.

128 Chan EK, Hardie R-A, Petersen DC, Beeson K, Bornman RM, Smith AB, Hayes VM: Revised timeline and distribution of the earliest diverged human maternal lineages in southern Africa. PloS one 2015, 10:e0121223.

129 Boattini A, Castr\’\i L, Sarno S, Useli A, Cioffi M, Sazzini M, Garagnani P, de Fanti S, Pettener D, Luiselli D: mtDNA variation in East Africa unravels the history of afro-asiatic groups. American journal of physical anthropology 2013, 150:375–385.

130 Rando J, Pinto F, Gonzalez A, Hernandez M, Larruga J, Cabrera V, Bandelt HJ: Mitochondrial DNA analysis of Northwest African populations reveals genetic exchanges with European, Near-Eastern, and sub-Saharan populations. Annals of human genetics 1998, 62:531–550.

131 Elmadawy MA, Nagai A, Gomaa GM, Hegazy HM, Shaaban FE, Bunai Y: Investigation of mtDNA control region sequences in an Egyptian population sample. Legal Medicine 2013, 15:338–341.

132 Mikkelsen M, Fendt L, Röck AW, Zimmermann B, Rockenbauer E, Hansen AJ, Parson W, Morling N: Forensic and phylogeographic characterisation of mtDNA lineages from Somalia. International journal of legal medicine 2012, 126:573.

133 Batai K, Babrowski KB, Arroyo JP, Kusimba CM, Williams SR: Mitochondrial DNA diversity in two ethnic groups in Southeastern Kenya: perspectives from the northeastern periphery of the Bantu expansion. American journal of physical anthropology 2013, 150:482–491.

134 Badro DA, Douaihy B, Haber M, Youhanna SC, Salloum A, Ghassibe-Sabbagh M, Johnsrud B, Khazen G, Matisoo-Smith E, Soria-Hernanz DF, others: Y-chromosome and mtDNA genetics reveal significant contrasts in affinities of modern Middle Eastern populations with European and African populations. PloS one 2013, 8:e54616.

135 Kivisild T, Reidla M, Metspalu E, Rosa A, Brehm A, Pennarun E, Parik J, Geberhiwot T, Usanga E, Villems R: Ethiopian mitochondrial DNA heritage: tracking gene flow across and around the gate of tears. Am. J. Hum. Genet. 2004, 75:752–70.

136 Rosenberg T, Preusser F, Fleitmann D, Schwalb A, Penkman K, Schmid T, Al-Shanti M, Kadi K, Matter A: Humid periods in southern Arabia: windows of opportunity for modern human dispersal. Geology 2011, 39:1115–1118.

137 Bekada A, Fregel R, Cabrera VM, Larruga JM, Pestano J, Benhamamouch S, González AM: Introducing the Algerian mitochondrial DNA and Y-chromosome profiles into the North African landscape. PLoS ONE 2013, 8:e56775.

138 Fernandes V, Triska P, Pereira JB, Alshamali F, Rito T, Machado A, Fajko\vsová Z, Cavadas B, \vCern\’y V, Soares P, others: Genetic stratigraphy of key demographic events in Arabia. PloS one 2015, 10:e0118625.

139 Poloni ES, Naciri Y, Bucho R, Niba R, Kervaire B, Excoffier L, Langaney A, Sanchez-Mazas A: Genetic evidence for complexity in ethnic differentiation and history in East Africa. Annals of human genetics 2009, 73:582–600.

140 Wood ET, Stover DA, Ehret C, Destro-Bisol G, Spedini G, McLeod H, Louie L, Bamshad M, Strassmann BI, Soodyall H, others: Contrasting patterns of Y chromosome and mtDNA variation in Africa: evidence for sex-biased demographic processes. European Journal of Human Genetics 2005, 13:867–876.

141 Vishnyatsky LB: The Paleolithic of Central Asia. Journal of World Prehistory 1999, 13:69–122.

142 Gaillard C, Singh M, Malassé AD: Late Pleistocene to early Holocene lithic industries in the southern fringes of the Himalaya. Quaternary International 2011, 229:112–122.

